# The environmental impact of pharmaceuticals: an evidence-mapping review of recent data on aquatic concentrations and predictable effects

**DOI:** 10.1101/2024.11.07.622417

**Authors:** Renda Francesca, Giunchi Valentina, Bianconi Matilde, Matera Rossana, Tandurella Emanuele, Poluzzi Elisabetta, Macedonio Giorgia, Lunghi Carlotta

## Abstract

Pharmaceuticals are recognised among emerging contaminants, particularly in water. They have the potential to alter ecosystem dynamics, with notable examples including hormone-induced feminization of male fish and disruptions to oogenesis in invertebrates. To assess the risk posed by pharmaceuticals, it is essential to understand their amount (via Measured Environmental Concentrations – MEC) and their actual effects on target species (via Predicted No Effect Concentrations – PNEC). Recently, many studies have aimed to collect MEC data from around the world, but a comprehensive overview is still lacking. Thus, the objective of this study is to provide a comprehensive overview by examining recently published literature on MEC data for a wide range of pharmaceuticals. Additionally, to enable risk assessment, this study also reviewed the published literature on PNEC data and integrated it with existing databases. A total of 315 substances were selected for MEC data extraction, with the inclusion of 56 articles. The most frequently monitored locations were Cadiz Bay in Spain (90 samples), the River Thames in the UK (51), and Hrdějovice in the Czech Republic (49). Most MEC samples were collected from surface water (N=325), influent wastewater treatment plants (WWTP) (205), and effluent WWTP (118). Based on PNEC values, risk analysis identified 81 pharmaceuticals as high-risk, with the highest risk values for propranolol (risk quotient [RQ]: 29,450,000), diclofenac (395,920), and 17alpha-ethinylestradiol (95,946). Additionally, the ATC classes with the most high-risk substances were anti-infectives (J), nervous system agents (N), cardiovascular agents (C), antineoplastic agents (L), analgesics (M), and sex hormones (G). The findings of this study highlight the widespread impact of pharmaceuticals across the globe and the involvement of multiple therapeutic classes. To move beyond the current point-in-time overview, which is limited to specific locations and sampling periods, systems for continuous monitoring of pharmaceuticals should be developed. This could involve the creation of resource-efficient methods and the integration of sampling data with estimation models. Furthermore, these results could serve as a starting point for developing and implementing actions to prevent and mitigate the environmental impact of pharmaceuticals.

## INTRODUCTION

Pharmaceuticals are increasingly recognised as a source of environmental pollution and are classified as emerging contaminants (Rivera-Utrilla et al. 2013). Their impact is widespread across different environmental matrices, with water being particularly affected (aus der Beek et al. 2016). This pollution originates from manufacture discharges, human and animal excretion via urine and feces, cleansing of the skin after applying topical treatments, and incorrect disposal of unused or expired medicines by flushing them down the drain (Peake et al. 2016). Pharmaceuticals in water systems can impact animal and plant population behaviours. Examples include hormone-induced feminization of male fish (Tyler and Jobling 2008), the development of antimicrobial resistance due to antibiotic exposure (Jones et al. 2023; Nhung et al. 2023), and growth and population issues caused by substances like analgesics (Gonzalez-Rey and Bebianno 2014; Parolini 2020). Even though the effects of some pharmaceuticals on aquatic flora and fauna are well-known, different geographic areas may be impacted in varying ways (Wilkinson et al. 2022). This variation can be attributed to differences in drug utilization patterns across regions, wastewater management and treatment practices, and characteristics of local stream networks. Additionally, since water passes through multiple stages—from influent wastewater to effluents, and from surface water to seawater—the effects of pharmaceuticals may differ at each stage (Pereira et al. 2020).

Furthermore, it is crucial to understand the actual risks posed by pharmaceutical concentrations. The standard measure used to assess the risk of pharmaceuticals in water involves the calculation of an active substance-specific (i.e., Active Pharmaceutical Ingredient (API)) risk quotient (RQ). This quotient is typically derived by comparing the Measured Environmental Concentration (MEC) to the Predicted No-Effect Concentration (PNEC) (FASS.se 2015). MEC represents the sampled concentration of the active substance in a water compartment, while PNEC represents the concentration limit below which adverse effects on organisms are not expected to occur.

The aim of this study is to review the published literature on water MEC to provide an overview of the recently sampled concentrations of substances, characterizing them by location and water type. Additionally, the study seeks to assess the risk posed by these by reviewing and retrieving related PNEC values. The results are presented according to the WHO-ATC (WHO) classification, offering insights into the therapeutic classes of greatest concern.

## METHODS

The present study focused on a group of pharmaceuticals previously identified by authoritative reports as posing potential ecological risks to the environment. Specifically, this work used as a starting point the following sources:

- The main European Union (EU) Directives on pharmaceutical pollution (Dir.2000/60/EC, Dir.2006/118/EC, Dir.2008/105/EC amended by Dir.2013/39/EC, Dir.2020/2184/CE) and the EU Commission Implementing Decision (2015/495, 2018/840, 2020/1161, and 2022/1307) establishing a Watch List of Substances for Union-wide monitoring in the field of water policy according to the European Parliament and Council Directive 2008/105/EC.
- EU and extra-EU (e.g., the United States and Australia) institutional projects.
- The report from the European Environmental Bureau (EBB), which was authored by a European environmental citizens’ organization in Germany (EEB -The European Environmental Bureau 2018).

The MEC and PNEC values of the extracted list of active substances were searched in the PubMed database with the following search string: “name of active substance” AND “environment” OR “disposal”. The search was performed for a period from January 1, 2019, to December 31, 2023, and only available open-access full-text articles were selected for review.

The active substances not included in the first list but present in the selected articles were also added to be screened. In this study, the selected articles contained MEC for at least one aquatic environment or PNEC for at least one aquatic organism. When multiple MEC values were reported from the same source, the highest value available was chosen to represent the worst-case scenario worldwide. Otherwise, if only aggregated data were reported, the mean value was used. Similarly, the lowest value was selected when multiple PNEC values were reported by the same source.

The environmental RQ for active substance with at least one MEC and one PNEC value was calculated as the ratio between MEC and PNEC. For substances where no PNEC was found in the literature, PNEC values were searched for in the NORMAN database (Dulio et al. 2020).

Based on the Swedish Association of Pharmaceutical Industries (FASS) criteria, environmental risks were classified into four categories. If the quotient resulted greater than 10, the risk was considered “high”. If it was between 10 and 1, the risk was classified as “moderate”. Similarly, if the quotient was between 1 and 0.1, it was labelled as “low”. Finally, the risk was deemed “insignificant” for quotients lower than 0.1(Lif 2012).

## Results

Screening EU directives and international projects on pharmaceuticals in the environment resulted in an initial selection of 86 active substances. Subsequent literature screening included 229 more substances.

The literature review of published MEC and PNEC data for the originally selected substances yielded 56 relevant articles (**Table 1**). Most of these articles were original research papers (N=33), while 8 were literature reviews. Some articles examined various contaminants (e.g., pharmaceuticals, pesticides, and personal care products), while others focused on specific therapeutic classes or substances. Studies reporting MEC values had either global-scale or specific-territory sampling (Table S2). For articles reporting PNEC values, tests were performed on various species but primarily on fish, algae, and crustaceans. Most of the data were related to surface water (Table S3).

**Table 1.**
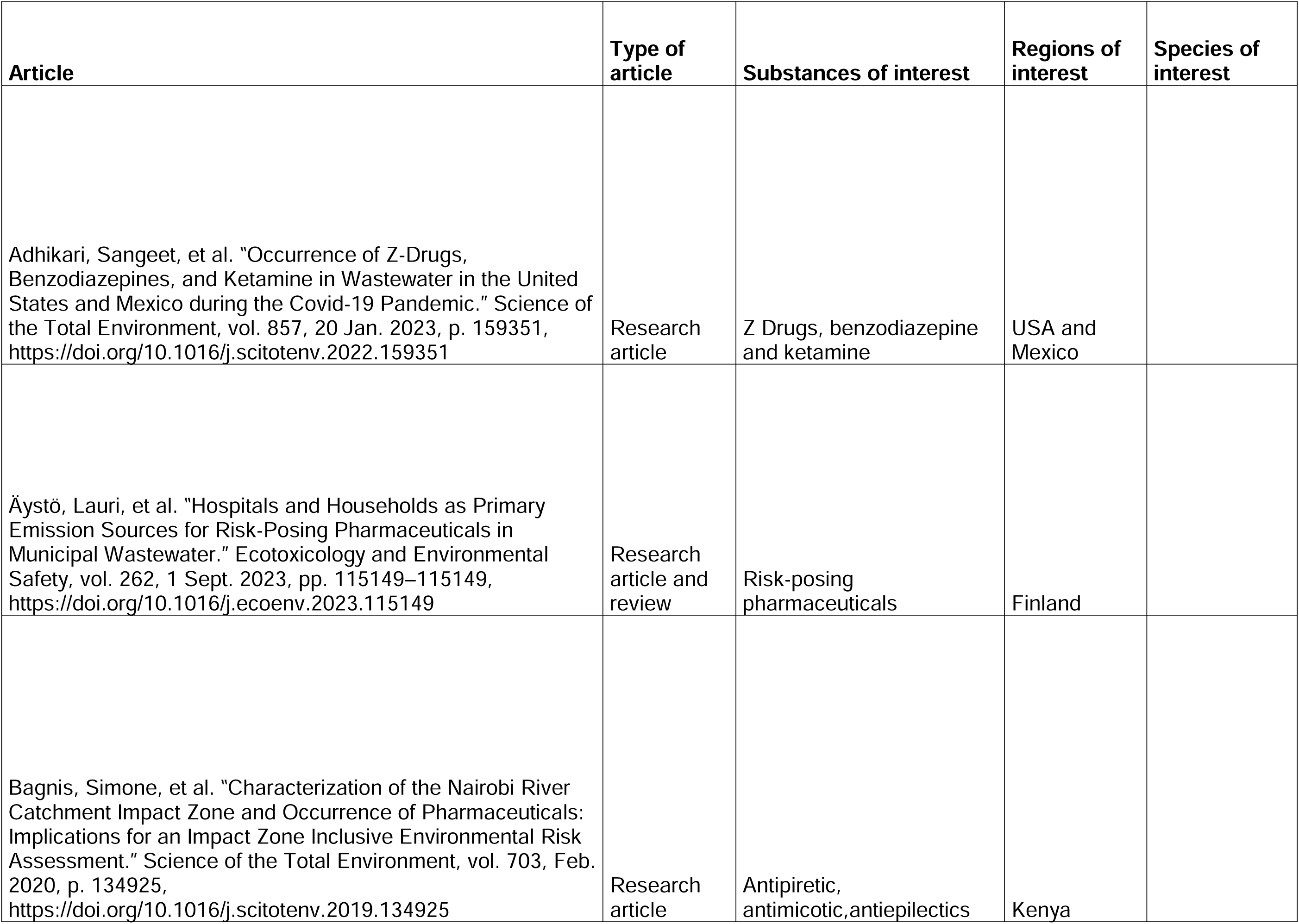
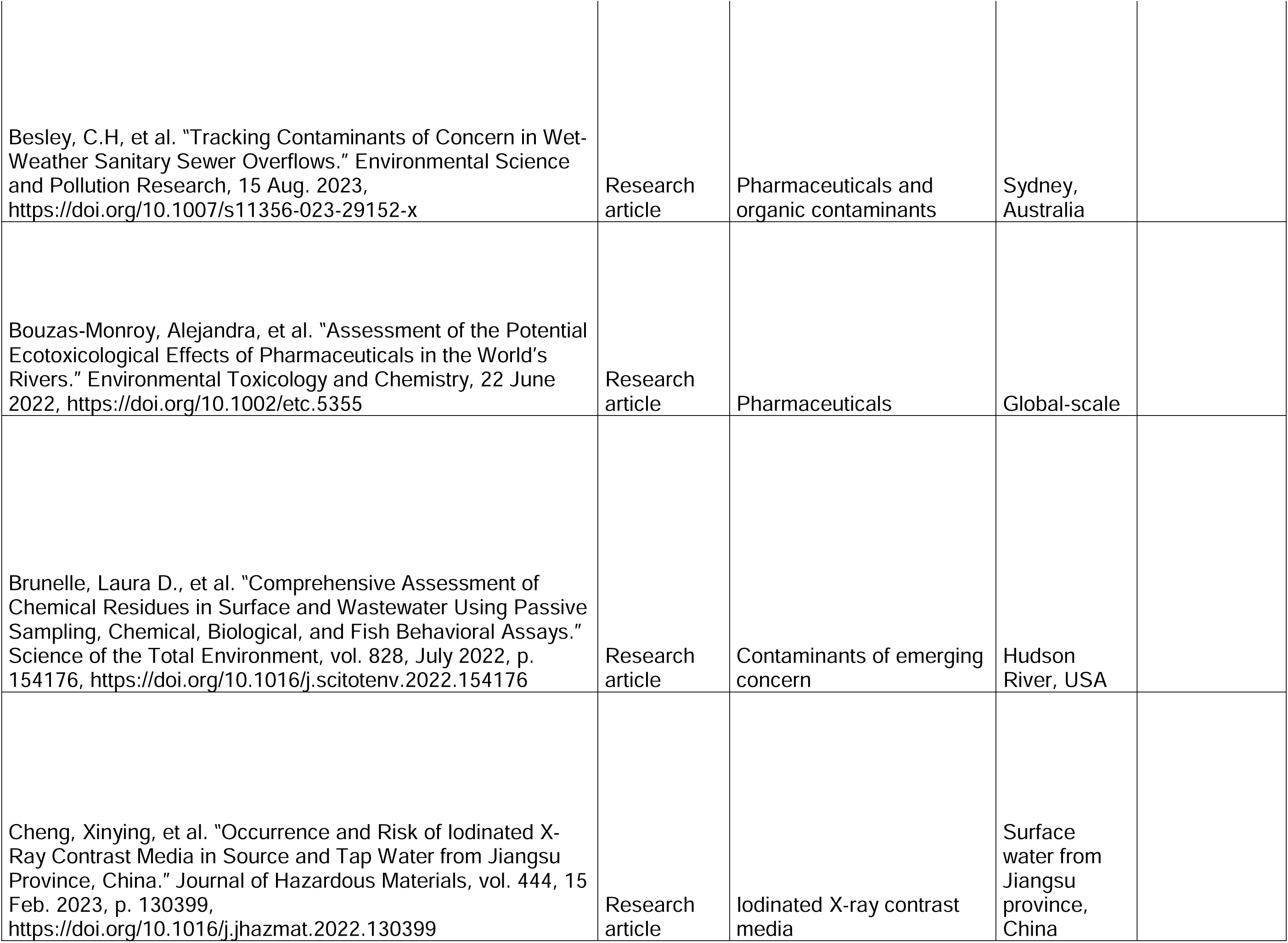
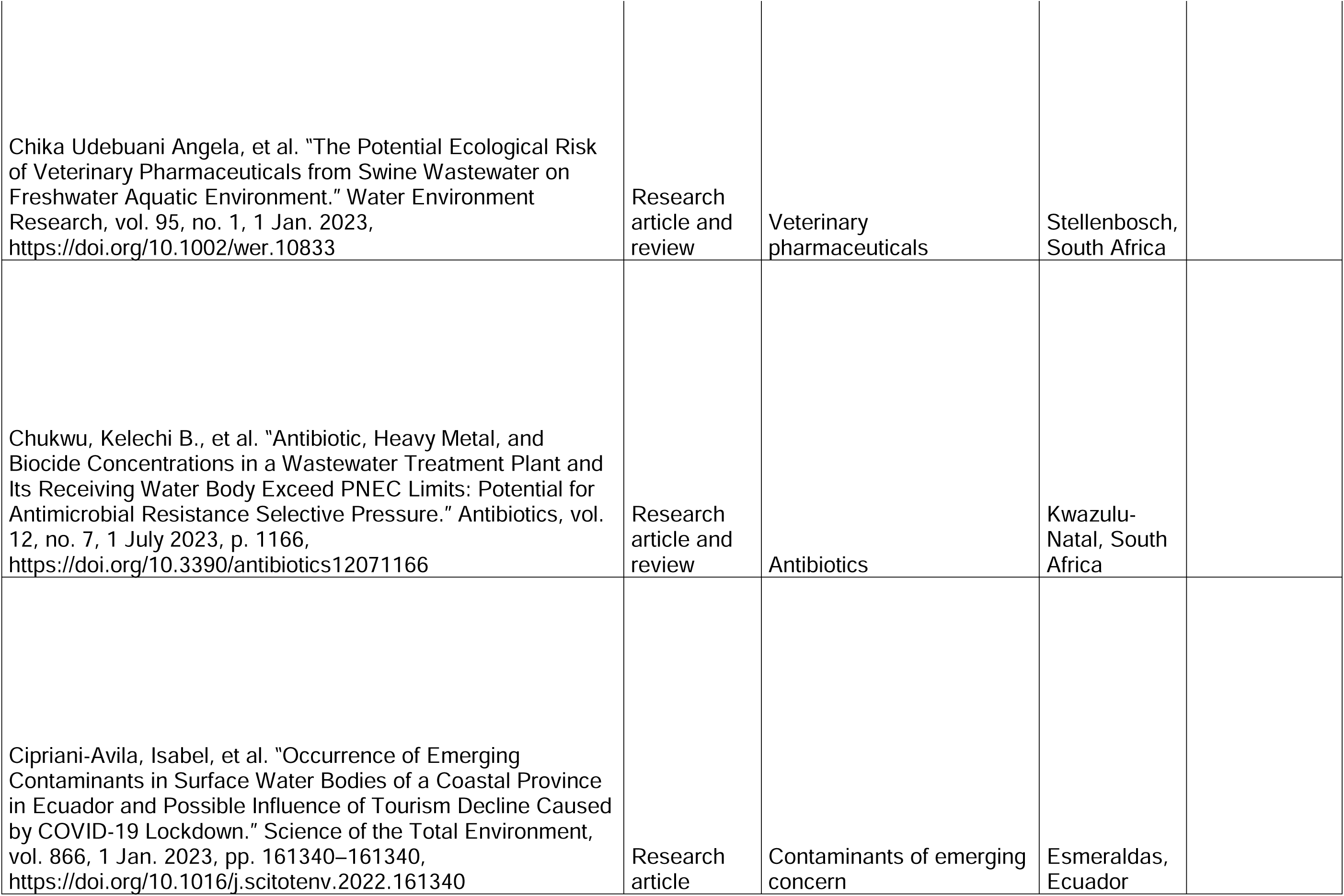
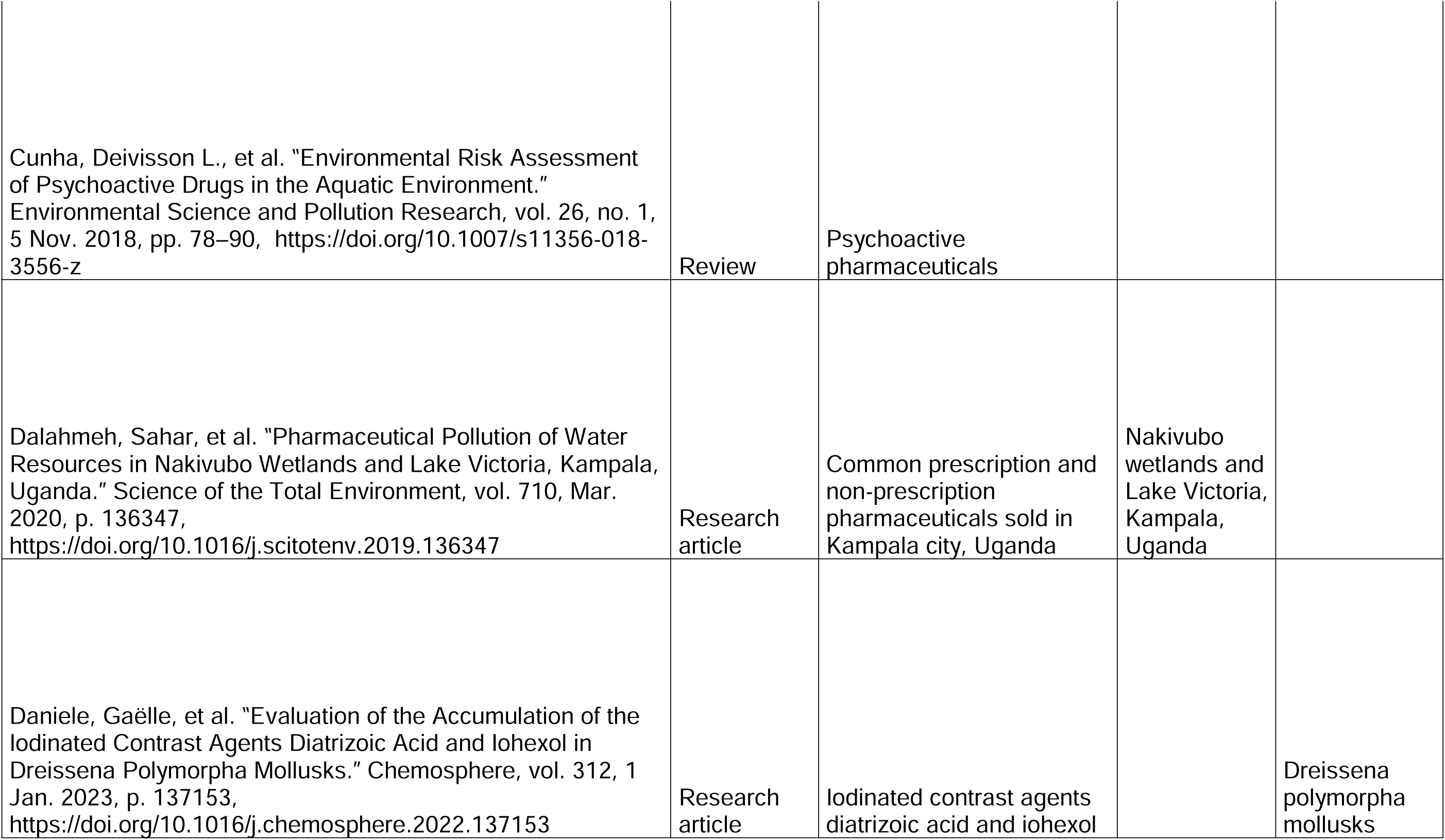
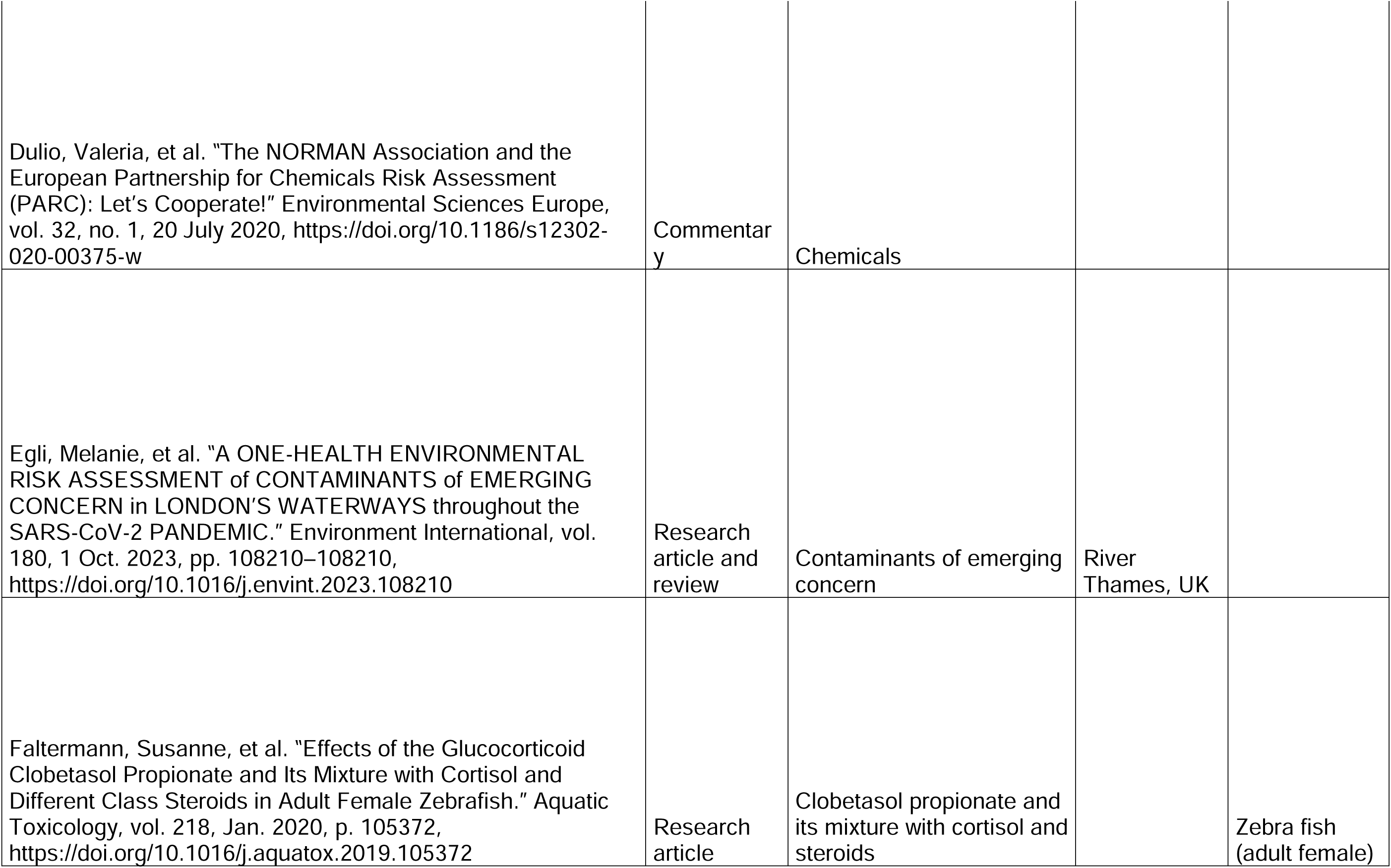
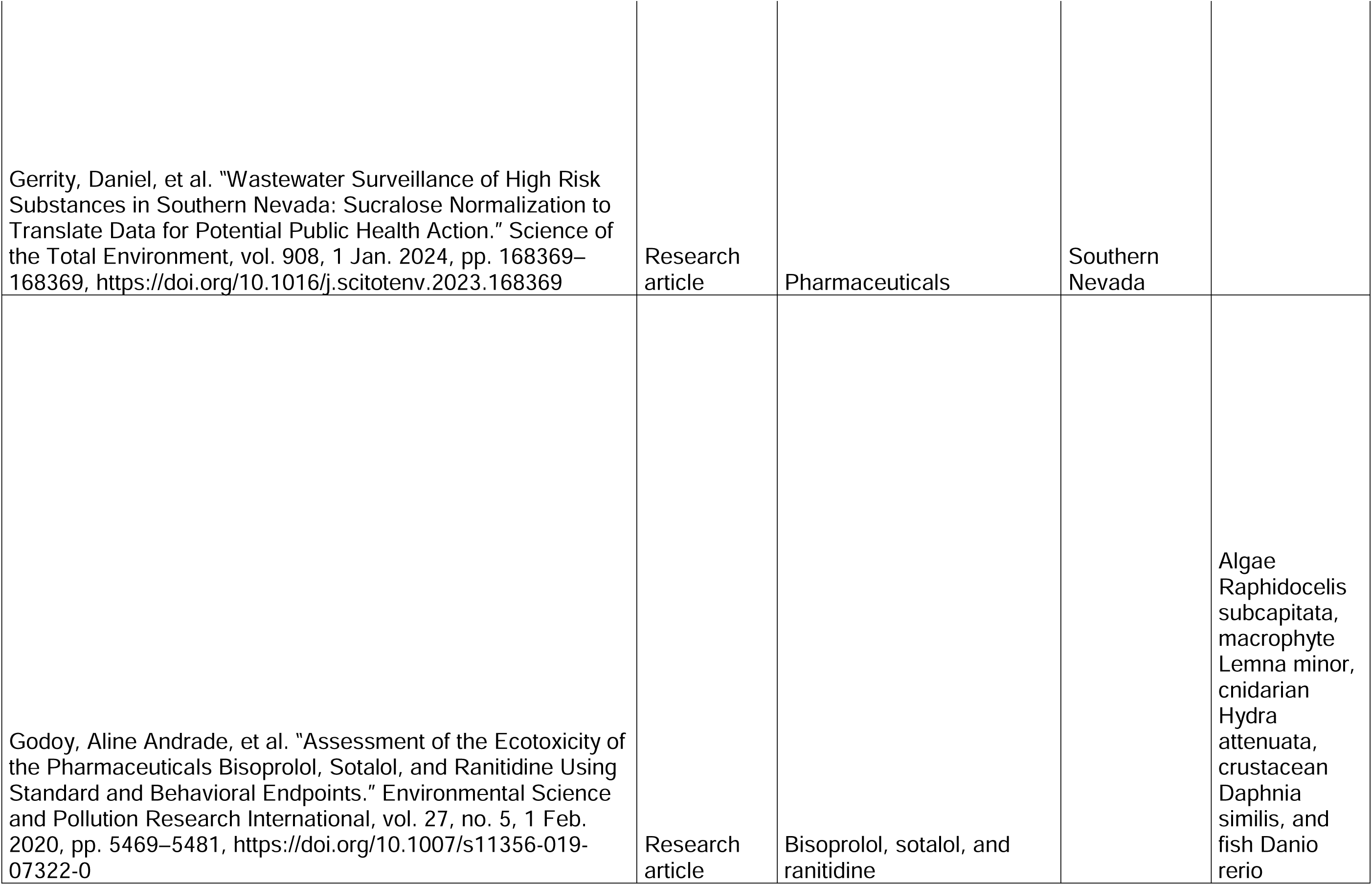
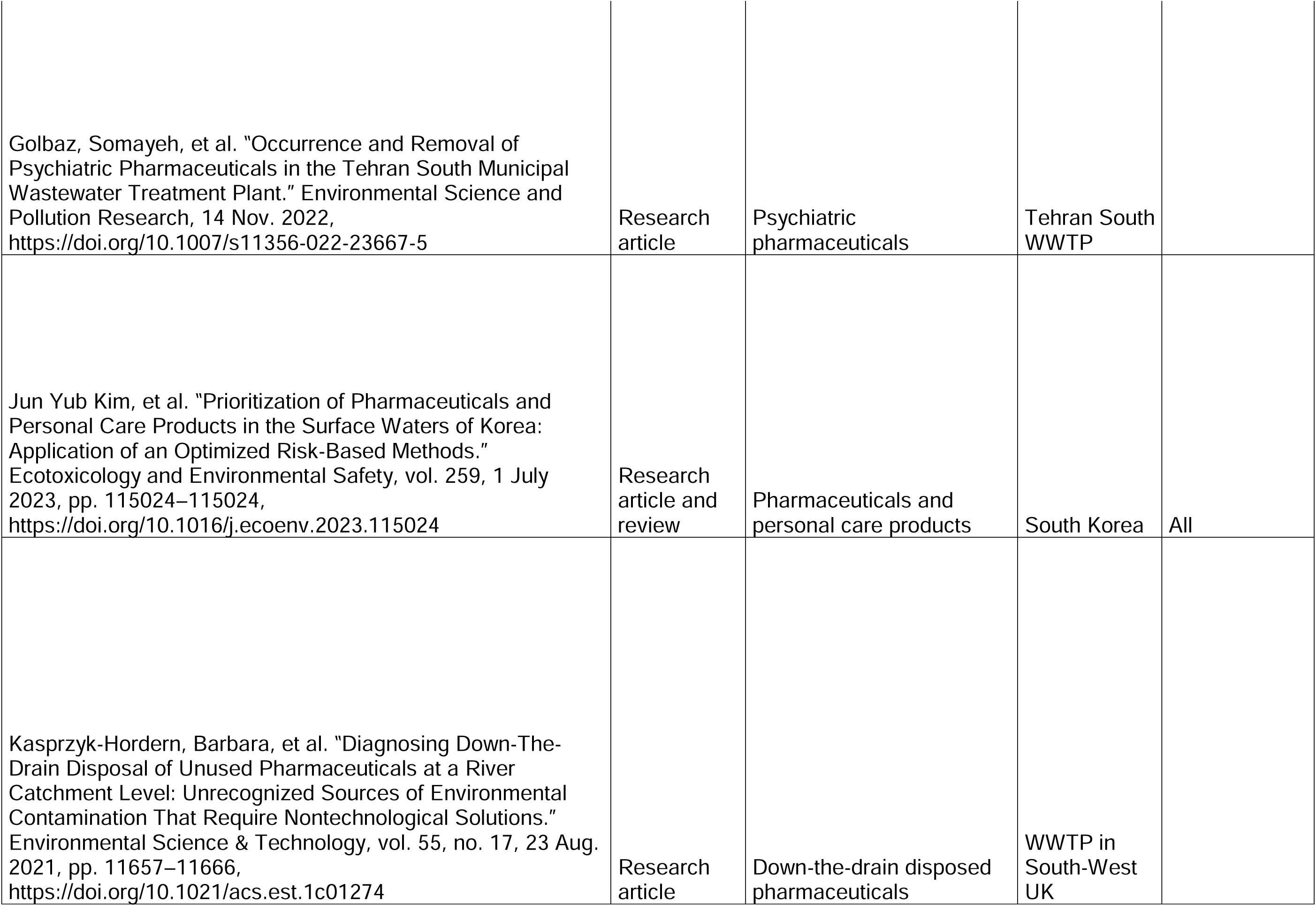
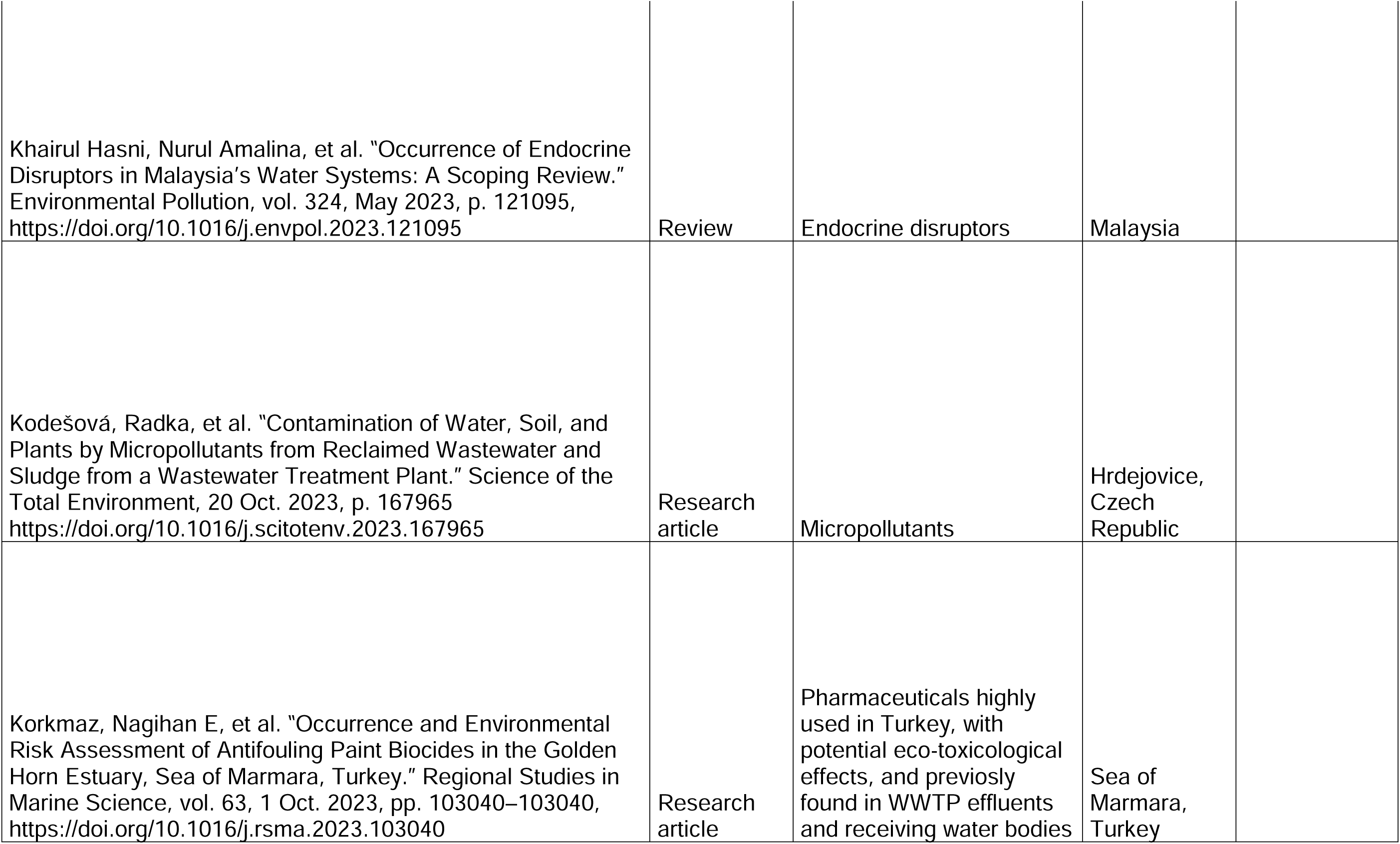
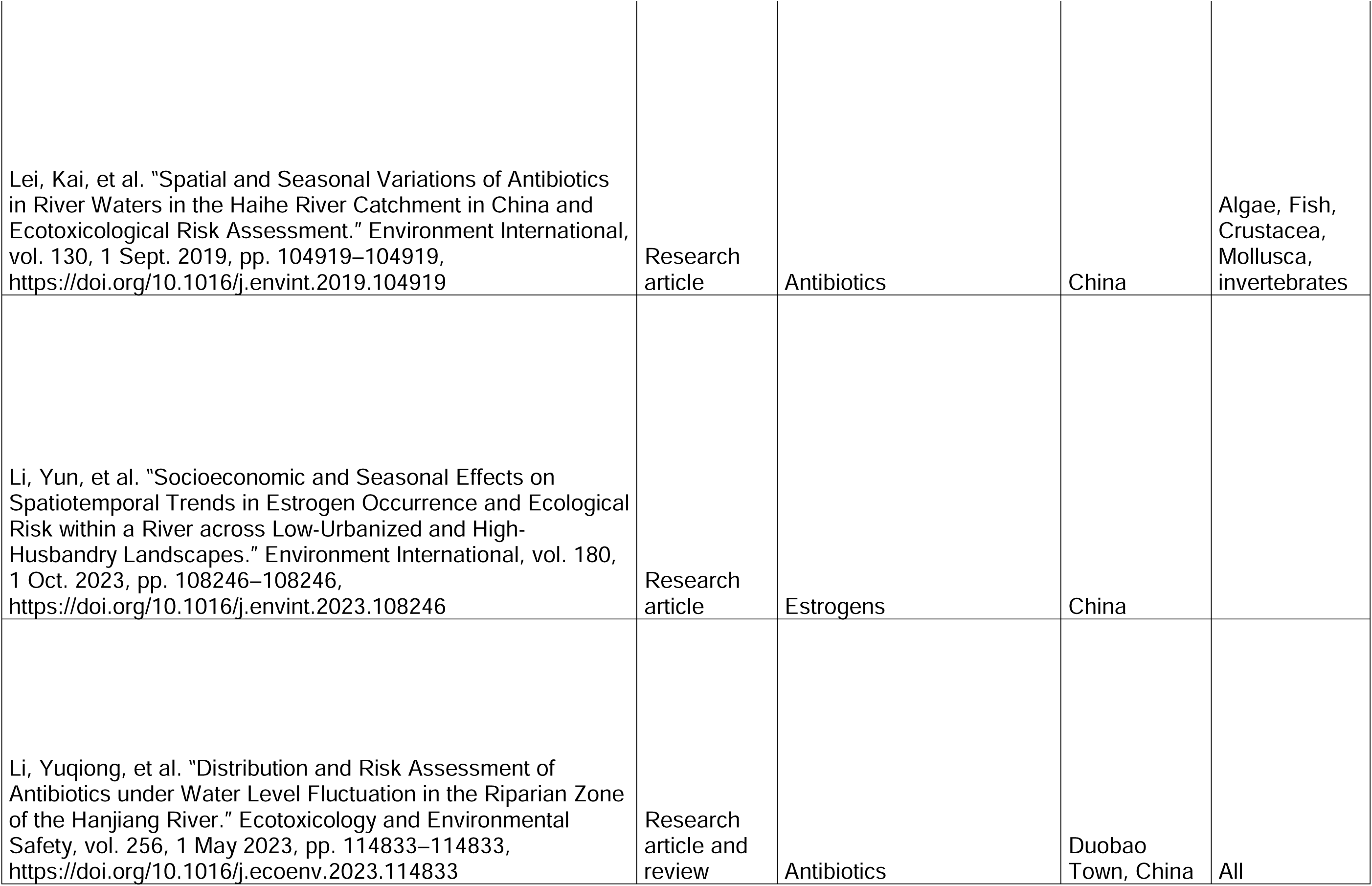
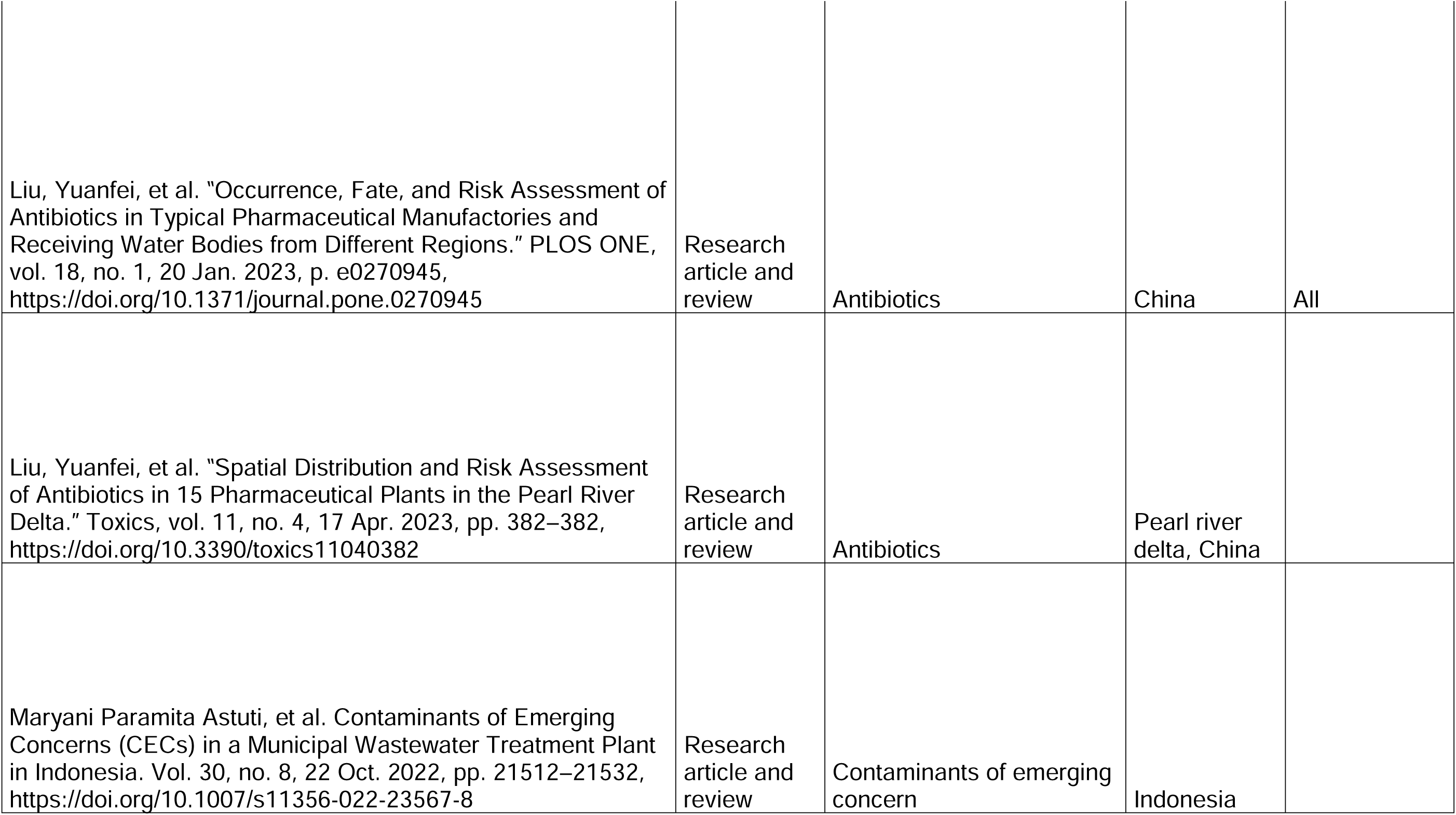
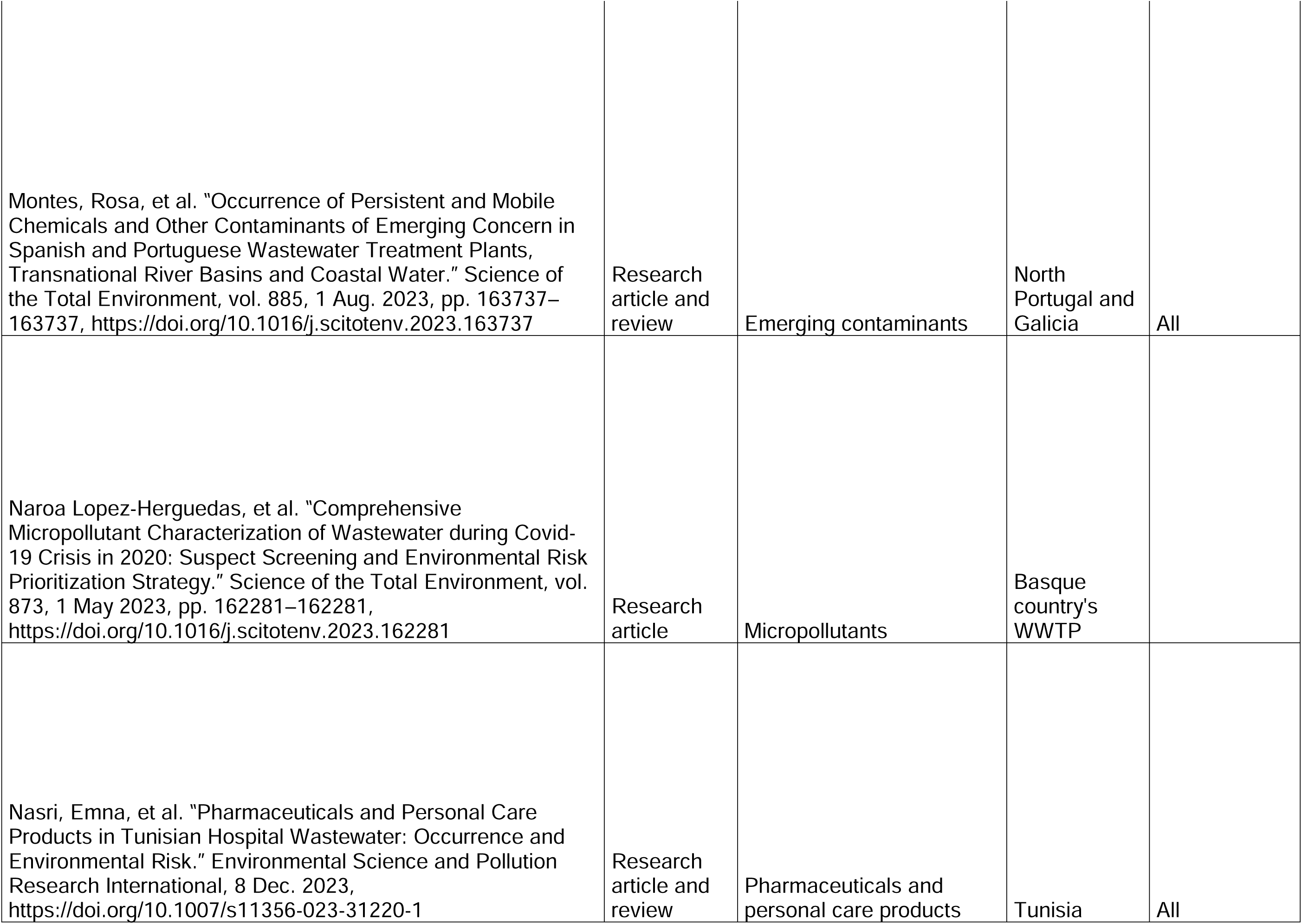
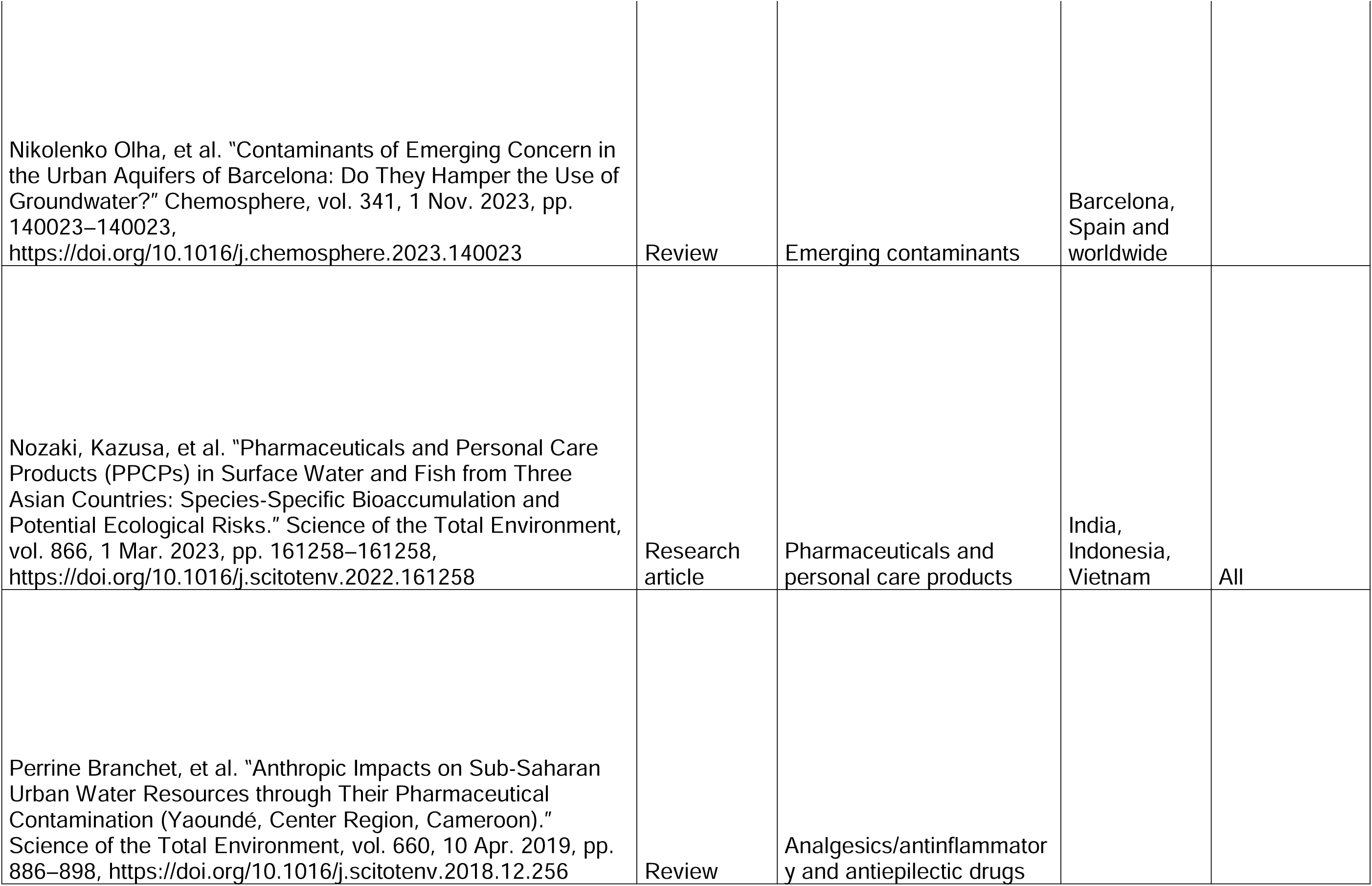
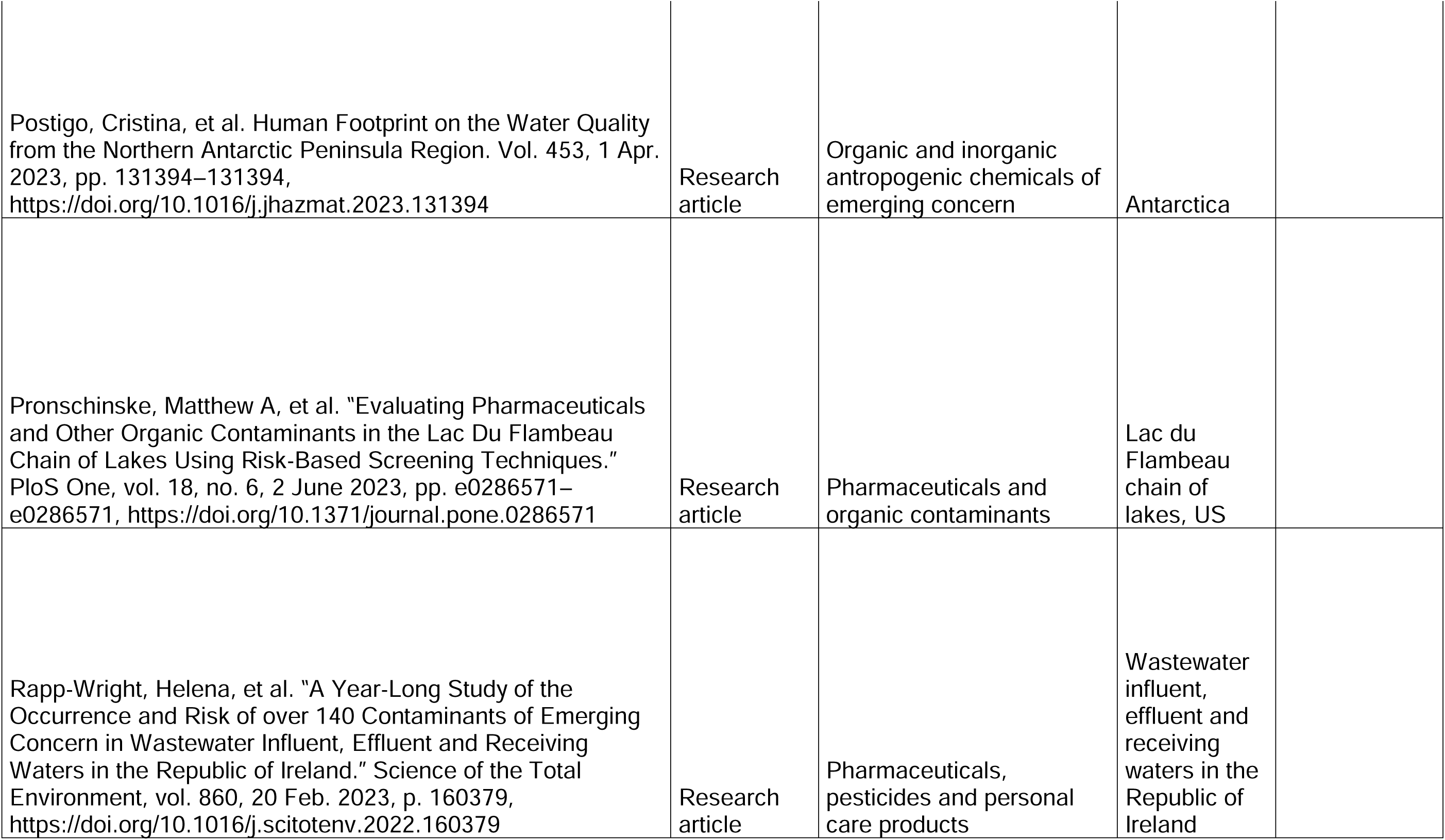
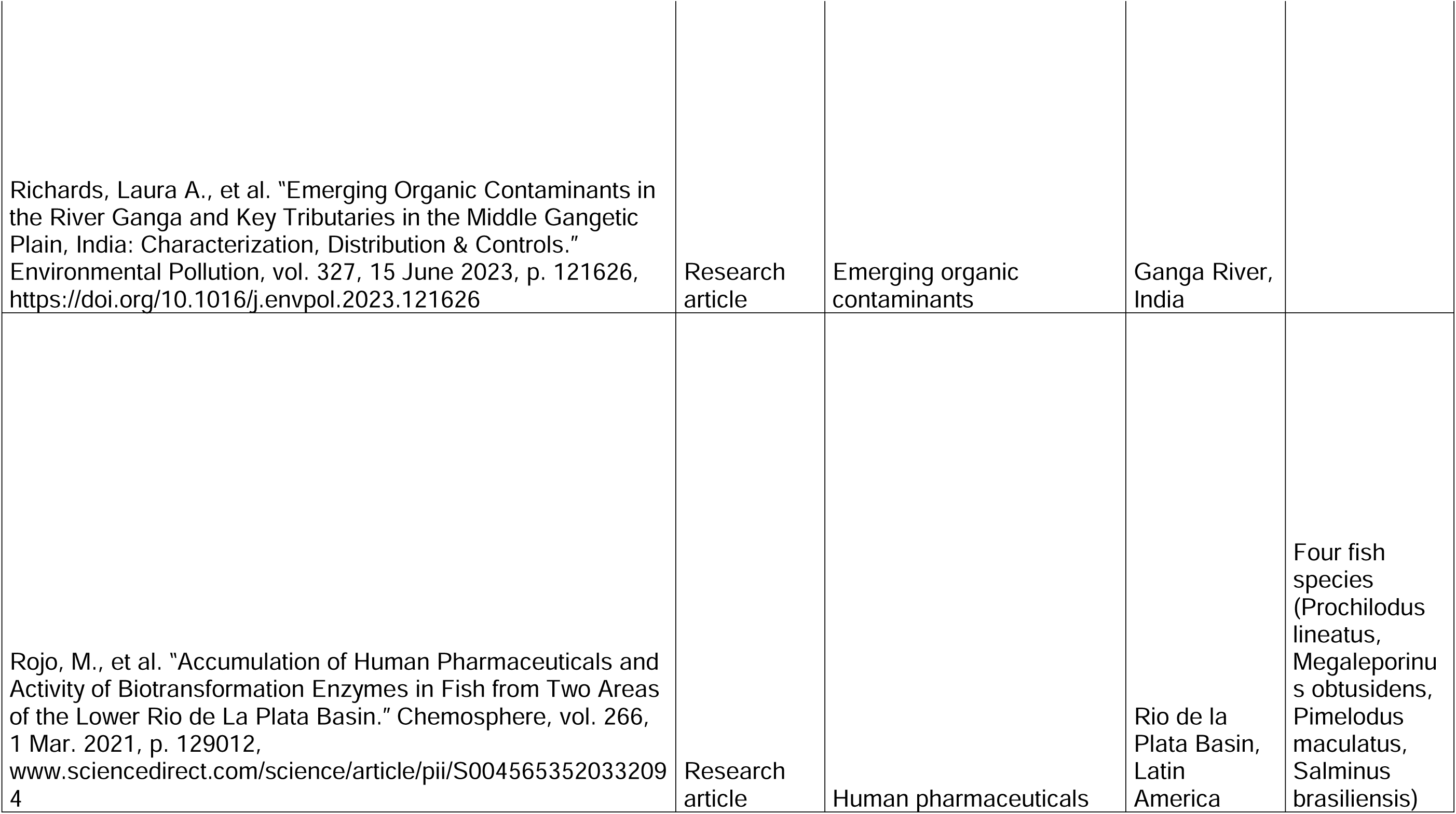
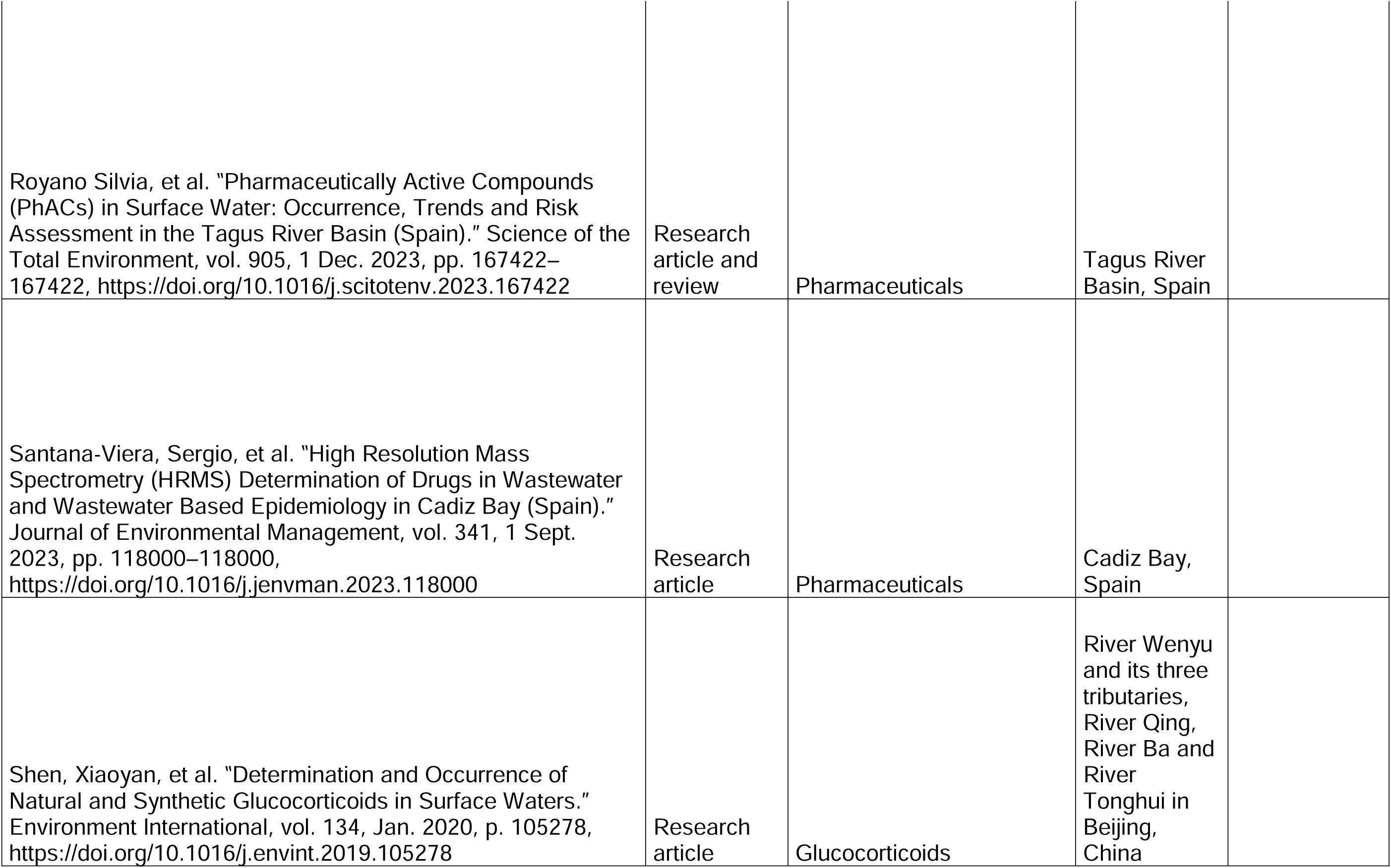
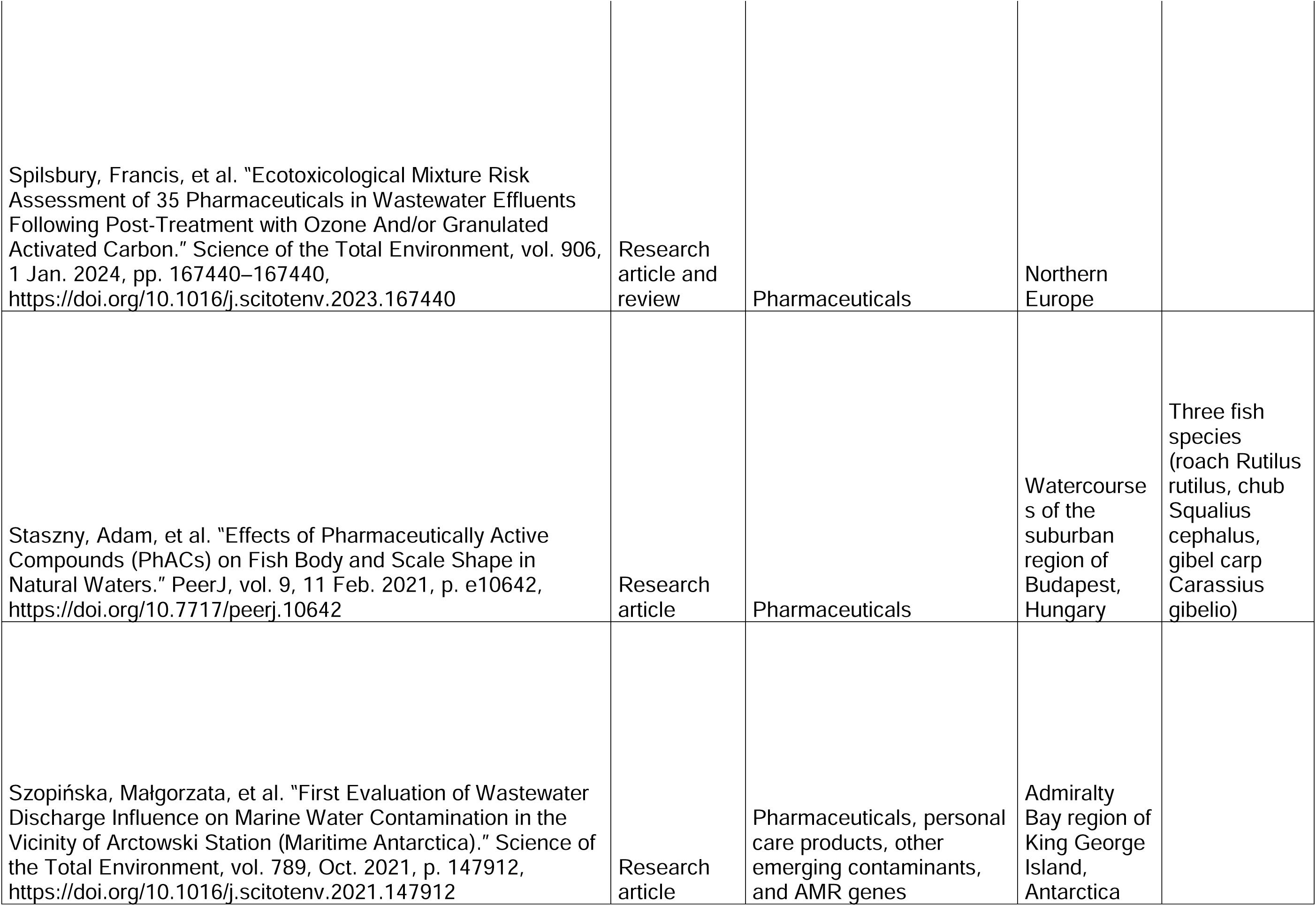
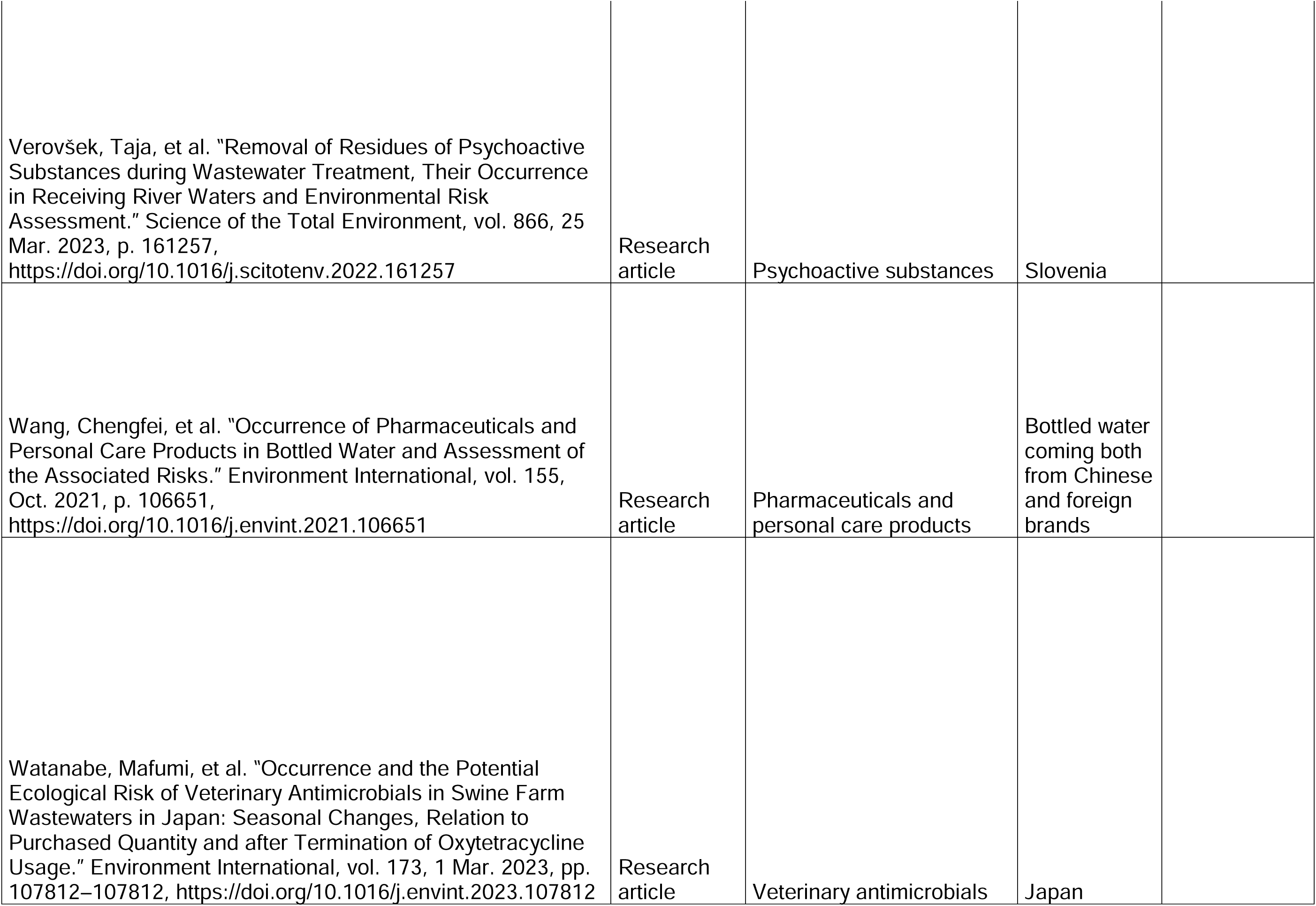
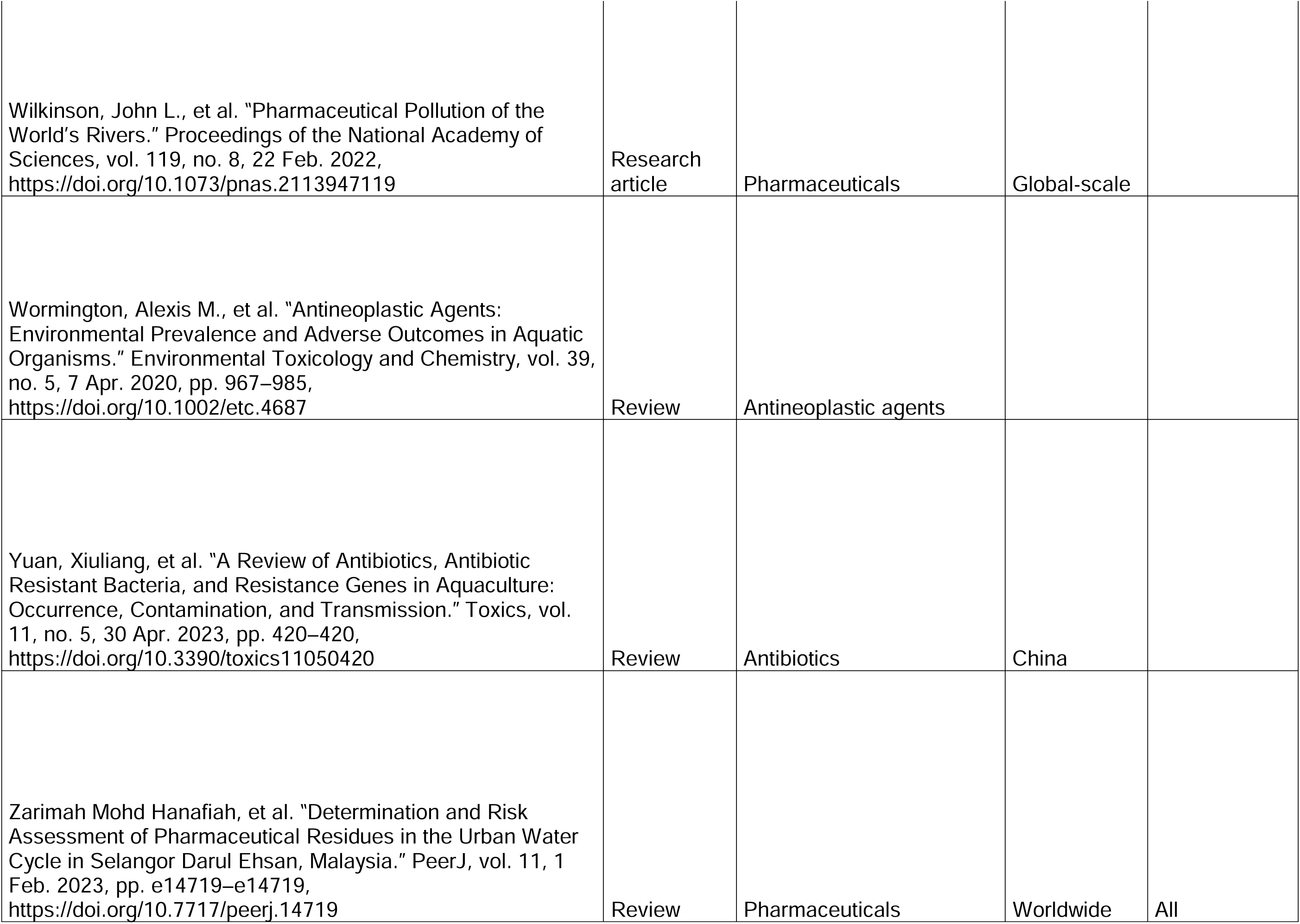
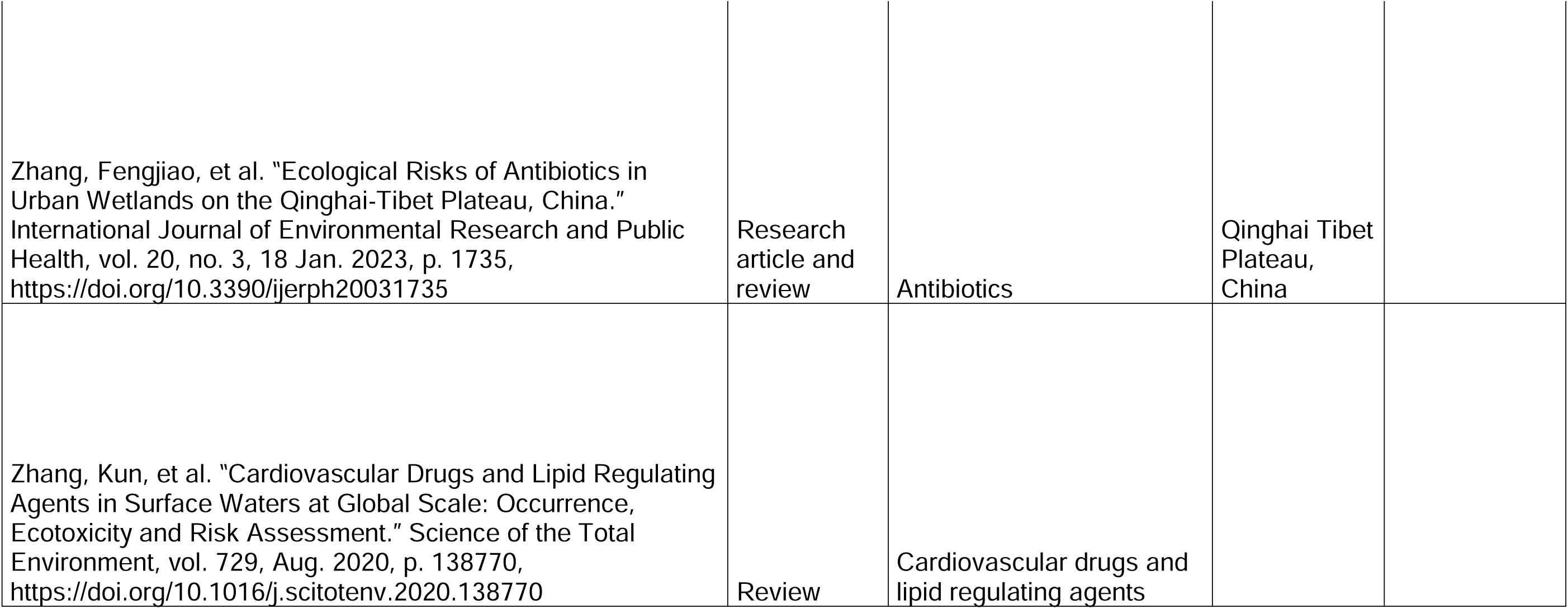
Articles selected from literature review and related information.

A total of 315 active substances with potential environmental risks belonging to 14 different drug classes (based on the anatomical therapeutic classification – ATC – 1^st^ level), including metabolites and veterinary drugs, were identified and selected. The MEC data were available for 314 of the 315 substances identified (**Table S1 and S2**). Notably, multiple MEC values from different articles were extracted for 162 active substances, including 24 for carbamazepine, 22 for trimethoprim, and 21 for sulfamethoxazole (**Table S2**). The highest MEC values were recorded for propranolol in wastewater in the United Kingdom (589,000,000 ng/L), paracetamol in hospital wastewater in Tunisia (1,260,000 ng/L) and sucralose in wastewater in the United States (1,100,000 ng/L).

The most monitored locations were Cadiz Bay in Spain (90 samplings), the river Thames in the UK (51 samplings), and Hrdějovice in the Czech Republic (49 samplings). The majority of MEC samplings were obtained from surface water (N=325), influent wastewater treatment plants (WWTP) (N=205), and effluent WWTP (N=118).

PNEC data were available for 217 of the 315 substances. Multiple PNEC values from various sources were extracted for 90 of these. Specifically, 9 values were retrieved for carbamazepine and trimethoprim, and 8 for diclofenac. The lowest PNEC values were recorded for 17alpha-ethinylestradiol in vertebrates (0.037 ng/L) and fish (0.44 ng/L), and for diclofenac in fish (0.05 ng/L) (**Table S3**).

The maximum environmental risk was calculated by comparing the highest retrieved MEC with the lowest retrieved PNEC. The analysis revealed that 81 active substances pose a high environmental risk (**Figure 1 and Table S1**). Of these, propranolol (RQ 29,450,000), diclofenac (RQ 395,920) and 17alpha-ethinylestradiol (RQ 95,946) were those with the highest environmental risk (Figure 1). The ATC classes with more active asubstances at potential high or moderate risk for the environment were anti-infectives for systemic use (J, with 30 at high risk and 9 at moderate risk); agents acting on the nervous system (N, with 12 at high risk and 10 at moderate risk), and cardiovascular system agents (C, with 9 at high risk and 5 at moderate risk). (**Table S1 and Figure 2**).

**Figure 1.**
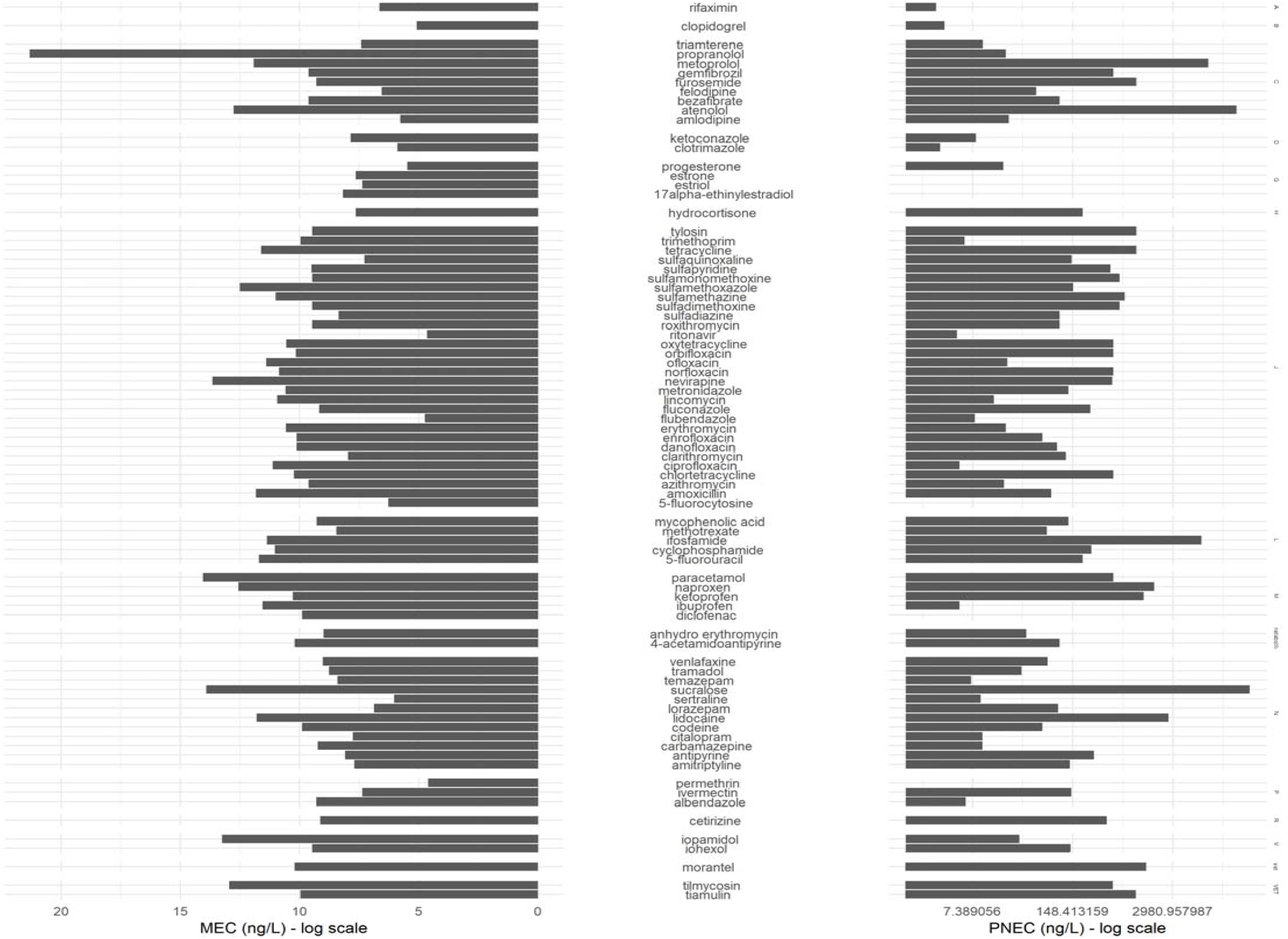
Bar plots of MEC (ng/L) and PNEC (ng/L) values for high-risk medicines, categorized by ATC class. Values are presented on a logarithmic scale.

**Figure 2:**
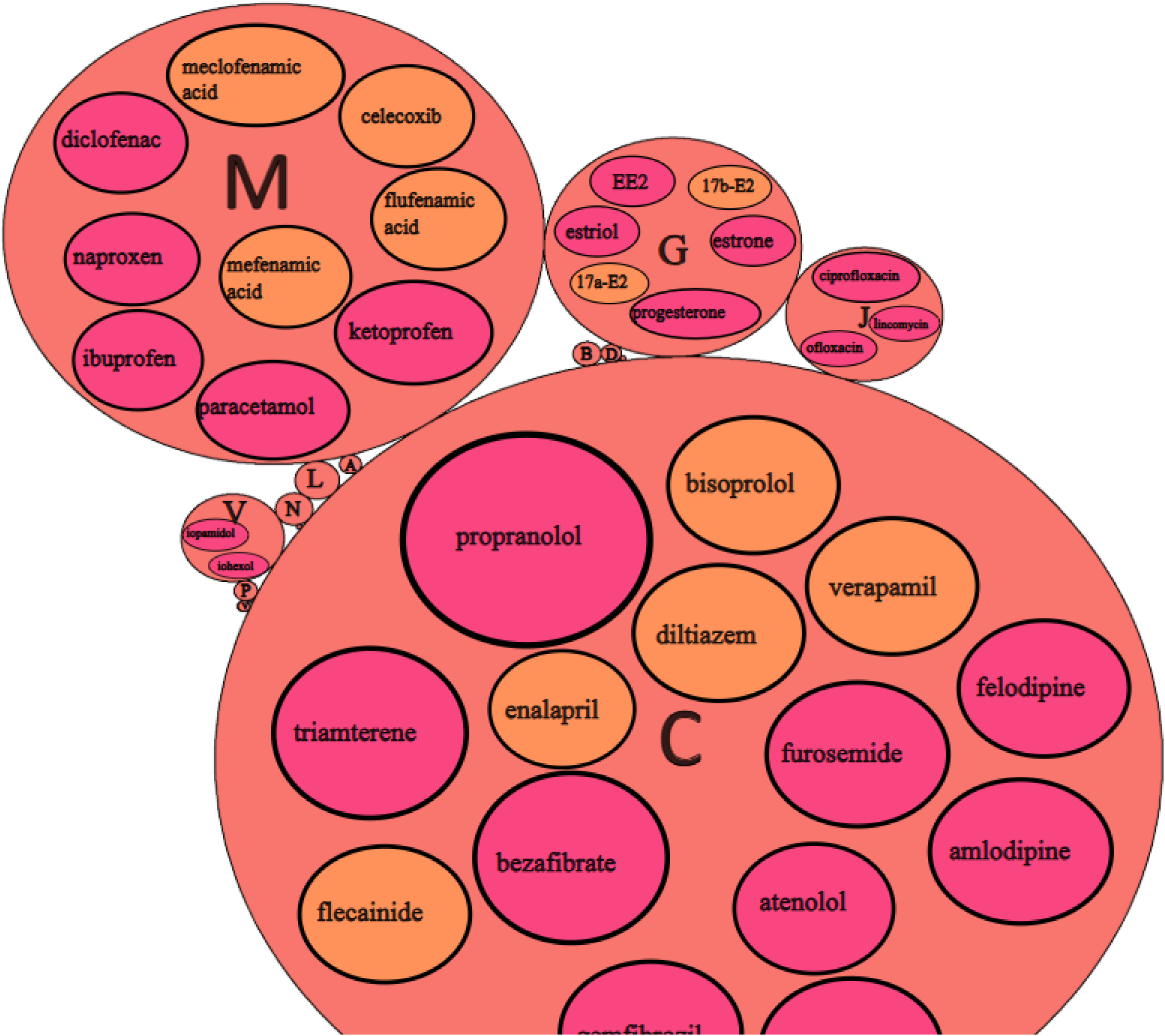
Circle packing chart illustrating the environmental risk of ATC classes and their medicines. The size of ATC circles is proportional to their risk level. Only higher-risk ATC classes and medicines are shown

Anti-infectives, antivirals, and antimycotics at high or moderate risk were found in different types of water across various continents. The highest values detected in Asia and Africa from surface water and effluent WWTP, respectively (**Table S2 and S3**). The nervous system agents posing a potential high or moderate risk to the environment were temazepam, citalopram, tramadol, codeine, carbamazepine, nicotine, and sertraline. The concentrations leading to these risks were from surface waters and WWTP influents and were detected in Western Europe and the Middle East (**Table S1**). The active substances belonging to ATC C that emerged at risk included two antihypertensives (propranolol and diltiazem), two lipid-lowering agents (gemfibrozil and bezafibrate), and the diuretic triamterene (**Figure 2** and **Tables S1**).

Other therapeutic classes with substances posing a risk to the environment included ATC L, where high-risk concentrations mainly originated from hospital wastewater, ATC M, which includes non-steroidal anti-inflammatory drugs (NSAIDs), ATC G, primarily estrogens used in contraception, ATC P consisting of anti-parasitic agents, and ATC R (**Figure** 2 and **Tables S1**).

## Discussion

This review article aimed to map the availability of recent and updated data on the MEC and PNEC of 315 active substances identified as potential environmental safety concerns and estimate their environmental risk. Among them, 81 were estimated at high environmental risk, and 40 at moderate risk. The ATC classification with the highest number of high-risk pharmaceuticals was class J, together with the topical intestinal antimicrobial rifaximin (ATC A). Among these, antibiotics in surface water are a primary concern. Indeed, they often exhibit toxicity to non-target organisms, particularly in aquatic environments, impacting primary producers and decomposers such as phytoplankton and zooplankton (Hillis et al. 2007; Ao et al. 2018). Antibiotics can also block the electron chain of photosystem II, increase oxidative stress (Willyard 2017; Kollef et al. 2017; García et al. 2020; Wang et al. 2021), and affect nutrient cycling, organic material degradation, sulphate reduction, and methanogenesis (Grenni et al. 2018; Xiong et al. 2019). Moreover, they tend to have broad-spectrum antibacterial activity, and their presence in surface water may contribute to microbial resistance. Indeed, microorganisms, including bacteria and fungi, develop resistance to antibacterial substances due to exposure to low concentrations over several generations. Microbial diversity plays a crucial role in biogeochemical cycles in soil and water, influencing various biological processes. Certain antibiotics have been demonstrated to affect the nitrogen cycle, resulting in heightened N_2_O release, diminished NO_3_ dissimilation, and a decline in nirZ and nirS gene potential in paddy fields (Shan et al. 2018). The risks posed by antibiotics are a concern for both medical and environmental agencies. The large amount of data collected in this review likely results from strict monitoring activities related to this therapeutic class.

ATC N represents another class containing high-risk pharmaceuticals. In particular, this class includes citalopram and sertraline, two Selective Serotonin Reuptake Inhibitors (SSRIs), whose adverse effects on aquatic species encompass weight loss, decreased offspring size, lower heart rate, and increased predation time (Stoczynski and van den Hurk 2020; Heyland et al. 2020; Ziegler et al. 2021). Moreover, tramadol and codeine, two opioids, have been observed to alter fish behavior.

Tramadol has been reported to slow the speed of fish and decrease their alertness, causing drowsiness, dizziness, loss of appetite, and impaired sensitivity to environmental stimuli (Langley et al. 2010; Buřič et al. 2018). Citalopram, on the other hand, has been reported to reduce aggressive and impulsive behaviors in fish, while also decrease the desire for reward (Tiihonen et al. 1996; Armenteros and Lewis 2002).

The third ATC class exhibiting the highest prevalence of high and moderate risk pharmaceuticals is class C. In particular, antihypertensives and lipid-lowering agents have been shown to affect the cardiac physiology, lipid metabolism, growth, and reproduction (e.g., spermatogenesis) of fish (Zhang et al. 2020).Among other pharmaceuticals identified as high or moderate risk are oncological treatments, belonging to ATC class L. These pharmaceuticals encompass cytotoxic and genotoxic substances that can have negative ecological impacts, induce DNA damage, disrupt endocrine pathways, reduce microbial diversity, persist through wastewater treatment processes, and bioaccumulate in aquatic organisms (A. Osawa et al. 2019; Russo et al. 2020). Their toxicity to aquatic organisms may be exacerbated by insufficient removal during wastewater treatment, resulting in their presence in groundwater globally (Ribeiro et al. 2022). Furthermore, high concentrations of anti-inflammatory and analgesic drugs (ATC M and N) in the environment can lead to serious ecological problems, including endocrine disruption, oxidative stress induction, and hormonal receptor binding in fish, invertebrates, and plants (Kwak et al. 2018; Silva et al. 2020; Parolini 2020; Svobodníková et al. 2020; Wijaya et al. 2020). Exposure to estrogens (ATC G) at high or moderate risk, even at low concentrations, can alter sex determination, decrease secondary sexual characteristics, and delay sexual maturity in affected organisms (Aris et al. 2014; Huang et al. 2015). Glucocorticoids (ATC H) have been shown to induce immunosuppression and reproductive disruption in fish (LaLone et al. 2012; Hidasi et al. 2017). Antiparasitic drugs (ATC P) impact numerous species of worms and insects involved in composting (Haseler et al. 2024), as is the case for antihistamines belonging to class R (Jonsson et al. 2015). Heterogeneity was observed in MEC data. While most of the available MEC data were from surface water, samples from other sources, such as influent and effluent WWTP, wastewater (unspecified, hospital, household, farm, manufacture), ground water, seawater, or bottled water were also included. Surface water may be the primary environment for many species, as it covers the largest areas and is thus most useful for understanding the actual risk. However, species also inhabit other aquatic environments, and all these environments are interconnected. Additionally, heterogeneity was evident among the sampling sites for the same substance and aquatic environment. For example, Africa and Asia emerge as high-risk areas with the highest concentrations of some antibiotics, while Europe and the Middle East have been found to have the highest concentrations of antidepressants. This could mirror different utilization patterns or wastewater treatment methods in different areas of the world, and it could be a starting point for future environmental and clinical sustainability improvements. Based on this prioritization list, which compiles sampling concentrations from around the world, targeted monitoring programs can be developed. Additionally, in areas that have not yet been subjected to monitoring, this list can serve as the basis for prioritizing monitoring efforts. While a substance’s exclusion from the selected list of pharmaceuticals does not necessarily indicate an absence of environmental risk, it is essential to acknowledge that emerging substances may not yet be incorporated into existing EU or international projects. Consequently, rigorous monitoring of new pharmaceuticals, especially throughout their life cycle, is crucial.

## Limitations

The main challenge faced during this review was the selection of active substances for inclusion. At present, there exists no comprehensive, accessible, and authoritatively recognized inventory of substances that may pose environmental concerns. The selection was based on European Union directives and international projects. To address the potentially limited number of active substances included in these sources, any substance with a potential environmental risk mentioned in any of the articles included in the literature search was incorporated. The data from the Region Stockholm database “Pharmaceuticals and the Environment” were excluded from the data sources due to the ongoing update of the PBT index (persistence, bioaccumulation, and toxicity information of pharmaceuticals), which rendered the extraction of uniform and comparable data problematic. It is also important to note that the literature review focused solely on the PubMed database and English language studies published in from 2019 to 2023 with open-access full text. Two articles, authored by Gerrity et al. and Spilsbury et al., respectively, despite being dated 2024, were made available online in the final months of 2023 and were consequently included. The selection of open-access publications may have limited the number of pharmaceuticals identified; however, this approach allows the dissemination of scientific knowledge at all levels of society with free and immediate access, adhering to principles of transparency and equity. Moreover, our review intentionally excluded non-therapeutic approved pharmaceuticals, such as illicit drugs, which may also impact aquatic environments. Another challenge is related to the attempt to link pharmaceuticals with the ATC classification. Indeed, some substances may correspond to more than one ATC code and, consequently, fall into multiple ATC category. In such cases, the most commonly used formulation of a pharmaceutical was prioritized, selecting the corresponding ATC code.

## Conclusions

Environmental pollution related to pharmaceuticals represents an emerging and concerning problem, and the presence of new sampling campaigns detecting these residues in previously understudied areas of the world (e.g., Africa and Asia) confirms its global nature. At the same time, research on the effects of pharmaceuticals in aquatic ecosystems is advancing, demonstrating adverse effects on animals and plants. In the long term, pharmaceuticals may exert detrimental impacts on the environment, especially through antimicrobial resistance, alteration of the endocrine system disruption, and growth inhibition. This review provides ana analysis of pharmaceuticals that potentially pose a significant risk to the environment, offering a global worst-case scenario risk assessment. Environmental risk has been identified across most regions of the world and across the majority of drug classes, with variations likely driven by differences in drug utilization patterns and wastewater management practices. Therapeutic areas with more substances identified at potential high or moderate risk to the environment included antimicrobials, agents acting on the nervous system, cardiovascular drugs, antineoplastic drugs, analgesics and sexual hormones. To move beyond the current point-in-time overview and enhance understanding the trends and behaviors of pharmaceutical concentrations in water, it is crucial to develop continuous monitoring systems. This development could encompass the creation of resource-efficient methods and the integration of sampling data with estimation models. Continuous monitoring could potentially inform actions toward more sustainable production, prescription, use, and disposal of medicines.

## Acknowledgements

F.R., R.M., E.T., M.B., and G.M. were not supported by any fund. V.G. and C.L. were supported by EU funds (*Programma Operativo Nazionale* Italian funds for green and innovative research based on European Structural and Investment Funds). E.P. was supported by institutional research funds (*Ricerca Fondamentale Orientata*).

## Statements and declarations

### Funding

The authors declare that no funds, grants, or other support were received during the preparation of this manuscript.

### Competing Interests

The authors have no relevant financial or non-financial interests to disclose.

### Author Contributions

F.R., R.M., E.T., and G.M. conceived and designed the study, and selected the pharmaceuticals to be included. All authors reviewed the articles included in the selection and extracted the relevant data. M.B. and V.G. examined the collected data, standardized them and analyzed them. The first draft of the manuscript was written by V.G., M.B., and F.R., and later, C.L. and E.P. commented on and integrated the previous versions of the manuscript. All the authors have read and approved the final manuscript.

## Notes

### Competing Interest Statement

The authors have declared no competing interest.

### Summary of Updates

The new version contains the change in the author's name order. Moreover, this include a new figure instead of the previous figure 1 and 2.

