## Supplementary material for "The environmental impact of pharmaceuticals: an evidence-mapping review of recent data on aquatic concentrations and predictable effects": Table S1 Table S2 Table S3: S1 - results with risk.pdf

| api | atc | mec_ng_l | pnec_ng_l | risk |
|---|---|---|---|---|
| rifaximin | A | 774 | 2,48 | 312,0967742 |
| ranitidine | A | 3350 | 3100 | 1,080645161 |
| metformin | A | 83111 | 160000 | 0,51944375 |
| vildagliptin | A | 455,4815 | 3018 | 0,150921637 |
| cimetidine | A | 3269,791 | 32090,49704 | 0,101892813 |
| pirenzepine | A | 13 | 170 | 0,076470588 |
| canagliflozin | A | 151 | 2856,074954 | 0,052869761 |
| sitagliptin | A | 3610 | 84000 | 0,04297619 |
| pioglitazone | A | 5 | 401 | 0,012468828 |
| omeprazole | A | 123 | 18100 | 0,00679558 |
| gliclazide | A | 13,682 | 2467 | 0,005546007 |
| tolbutamide | A | 5 | 1388 | 0,003602305 |
| acetohexamide | A | 5 | 4524 | 0,001105217 |
| clopidogrel | B | 162 | 3,21 | 50,46728972 |
| warfarin | B | 7000 | 11000 | 0,636363636 |
| ticlopidine | B | 69,687 | 421 | 0,165527316 |
| propranolol | C | 1800000000 | 20 | 90000000 |
| triamterene | C | 1660 | 10,02 | 165,6686627 |
| bezafibrate | C | 15100 | 100 | 151 |
| gemfibrozil | C | 15000 | 500 | 30 |
| atenolol | C | 350000 | 20000 | 17,5 |
| metoprolol | C | 150161 | 8600 | 17,4605814 |
| amlodipine | C | 322 | 22 | 14,63636364 |
| felodipine | C | 700 | 50 | 14 |
| furosemide | C | 11000 | 1000 | 11 |
| verapamil | C | 431 | 48 | 8,979166667 |
| diltiazem | C | 643 | 225,21 | 2,855113006 |
| bisoprolol | C | 5658 | 3150 | 1,796190476 |
| flecainide | C | 756 | 640,49 | 1,180346297 |
| enalapril | C | 1700 | 1575,29 | 1,079166376 |
| sotalol | C | 2000 | 2022 | 0,989119683 |
| olmesartan | C | 1090,2325 | 1404 | 0,776518875 |
| rosuvastatin | C | 719 | 1800 | 0,399444444 |
| chlorthalidone | C | 311 | 885,96 | 0,351031649 |
| dopamine | C | 15673 | 50305,97 | 0,31155348 |
| hydrochlorothiazide | C | 8900 | 34000 | 0,261764706 |
| losartan | C | 4359 | 27900 | 0,156236559 |
| atorvastatin | C | 1283 | 8500 | 0,150941176 |
| pentoxifylline | C | 693 | 6026,08 | 0,115000133 |
| eprosartan | C | 10449 | 100000 | 0,10449 |
| timolol | C | 25 | 310,281 | 0,080572127 |
| fenofibrate | C | 390 | 4900 | 0,079591837 |
| pravastatin | C | 354 | 4566,17 | 0,07752668 |
| telmisartan | C | 2739,4105 | 49000 | 0,055906337 |
| valsartan | C | 12737 | 560000 | 0,022744643 |
| valsartan acid | C | 4517 | 320000 | 0,014115625 |
| irbesartan | C | 5318 | 700000 | 0,007597143 |
| alprenolol | C | 10 | 1335 | 0,007490637 |
| candesartan | C | 448 | 100000 | 0,00448 |

| | | | | |
|---|---|---|---|---|
| mexiletine | C | 22 | 8460 | 0,002600473 |
| aliskiren | C | 213 | 1000000 | 0,000213 |
| ketoconazole | D | 2600 | 8,14 | 319,4103194 |
| clotrimazole | D | 362,215 | 2,8 | 129,3625 |
| triclosan | D | 515 | 110 | 4,681818182 |
| griseofulvin | D | 1,2 | 148,01 | 0,00810756 |
| climbazole | D | 203 | | |
| 17alpha-ethinylestradiol | G | 3550 | 0,037 | 95945,94595 |
| estriol | G | 1578 | 0,465 | 3393,548387 |
| estrone | G | 2090 | 1 | 2090 |
| progesterone | G | 240 | 18,6 | 12,90322581 |
| 17-beta estradiol | G | 3890 | 870 | 4,471264368 |
| 17alpha-estradiol | G | 8,491 | 2 | 4,2455 |
| norethisterone | G | 35,7 | 37 | 0,964864865 |
| testosterone | G | 468 | 4365,767 | 0,107197659 |
| medroxyprogesterone | G | 489 | 6723,61 | 0,072728787 |
| diethylstilbestrol | G | 3,62 | 64,27 | 0,056324879 |
| tamsulosin | G | 18 | 350 | 0,051428571 |
| finasteride | G | 46 | 5000 | 0,0092 |
| estradiol | G | 20,87 | | |
| hydrocortisone | H | 2100 | 200 | 10,5 |
| corticosterone | H | 1321 | 27951,65 | 0,04726018 |
| cortisone | H | 433 | 28118,44 | 0,015399147 |
| prednisolone | H | 94 | 24373,3 | 0,003856679 |
| ciprofloxacin | J | 68000 | 5 | 13600 |
| ofloxacin | J | 89600 | 21 | 4266,666667 |
| lincomycin | J | 56258,34 | 14 | 4018,452857 |
| trimethoprim | J | 21000 | 5,8 | 3620,689655 |
| 5-fluorocytosine | J | 531 | 0,22168 | 2395,344641 |
| erythromycin | J | 38952,31 | 20 | 1947,6155 |
| sulfamethoxazole | J | 268100 | 150 | 1787,333333 |
| nevirapine | J | 853556 | 481,82 | 1771,524636 |
| amoxicillin | J | 136380 | 78 | 1748,461538 |
| azithromycin | J | 15148 | 19 | 797,2631579 |
| enrofloxacin | J | 25000 | 60 | 416,6666667 |
| metronidazole | J | 39900 | 130 | 306,9230769 |
| danofloxacin | J | 25000 | 93,08 | 268,5861624 |
| roxithromycin | J | 12895,52 | 100 | 128,9552 |
| tetracycline | J | 110000 | 1000 | 110 |
| norfloxacin | J | 52301,91 | 500 | 104,60382 |
| sulfamethazine | J | 59825,12 | 700 | 85,46445714 |
| oxytetracycline | J | 38456,22 | 500 | 76,91244 |
| chlortetracycline | J | 28000 | 500 | 56 |
| orbifloxacin | J | 26000 | 500 | 52 |
| sulfadiazine | J | 4269,31 | 100 | 42,6931 |
| fluconazole | J | 9745 | 250 | 38,98 |
| sulfapyridine | J | 13310 | 460 | 28,93478261 |
| clarithromycin | J | 2900 | 120 | 24,16666667 |
| ritonavir | J | 105 | 4,61 | 22,77657267 |
| sulfadimethoxine | J | 13000 | 600 | 21,66666667 |

| | | | | |
|---|---|---|---|---|
| sulfamonomethoxine | J | 13000 | 600 | 21,66666667 |
| flubendazole | J | 114,947 | 7,90569415 | 14,53977321 |
| tylosin | J | 13000 | 1000 | 13 |
| sulfaquinoxaline | J | 1452,3 | 143,05 | 10,15239427 |
| ampicillin | J | 92,075 | 12 | 7,672916667 |
| sulfachlopyridazine | J | 5968,32 | 800 | 7,4604 |
| pefloxacin | J | 18569,23 | 2575,27 | 7,210595394 |
| doxycycline | J | 14000 | 2000 | 7 |
| sulfacetamide | J | 56931 | 14307,8 | 3,979018437 |
| itraconazole | J | 41,3 | 12,88 | 3,206521739 |
| sulfametoxydiazine | J | 2489,52 | 800 | 3,1119 |
| chloramphenicol | J | 153 | 130 | 1,176923077 |
| penicillin | J | 2200 | 2135,24 | 1,030329143 |
| marbofloxacin | J | 6300 | 7780 | 0,809768638 |
| efavirenz | J | 74 | 200,79 | 0,36854425 |
| sulfathiazole | J | 2875,69 | 11000 | 0,261426364 |
| streptomycin | J | 100 | 482,8664412 | 0,207096604 |
| sulfamerazine | J | 160 | 780 | 0,205128205 |
| penicillin v | J | 34 | 180,681 | 0,188176953 |
| sulfamethizole | J | 242 | 1495,22 | 0,161849092 |
| clindamycin | J | 296 | 1900 | 0,155789474 |
| enoxacin | J | 244 | 2514,2 | 0,097048763 |
| lopinavir | J | 121 | 1575 | 0,076825397 |
| flumequine | J | 114 | 1500 | 0,076 |
| oxolinic acid | J | 24 | 380 | 0,063157895 |
| clevudine | J | 26,331 | 1592 | 0,016539573 |
| cefadroxil | J | 28,465 | 2670 | 0,010661049 |
| fleroxacin | J | 11,12 | 1112,28 | 0,009997483 |
| lomefloxacin | J | 7,74 | 826,18 | 0,009368419 |
| cephalexin | J | 22,32 | 4026,24 | 0,005543634 |
| darunavir | J | 9,246 | 1815 | 0,005094215 |
| metacycline | J | 31,76 | 6270 | 0,005065391 |
| cefradine | J | 20,8815 | 5893,697436 | 0,003543022 |
| oseltamivir | J | 8,947 | 3352 | 0,002669153 |
| sofosbuvir | J | 5 | 3117 | 0,001604107 |
| sulfabenzamide | J | 4 | 3052,62 | 0,00131035 |
| atazanavir | J | 2 | 1771 | 0,001129305 |
| sulfamethoxypiridazine | J | 0,3 | 3820 | 7,8534E-05 |
| difloxacin | J | 16,14 | 240000 | 0,00006725 |
| sulfanoxaline | J | 0,8 | 19700 | 4,06091E-05 |
| sulfamidine | J | 9780 | | |
| sulfadoxine | J | 5789,31 | | |
| pipedemic acid | J | 574 | | |
| sulfamidics | J | 6,38 | | |
| 5-fluorouracil | L | 122000 | 200 | 610 |
| cyclophosphamide | L | 61660 | 260 | 237,1538462 |
| mycophenolic acid | L | 10739 | 130 | 82,60769231 |
| methotrexate | L | 4700 | 68,65 | 68,46321923 |
| ifosfamide | L | 86200 | 6963,88 | 12,378157 |
| bicalutamide | L | 2969 | 590 | 5,03220339 |

| | | | | |
|---|---|---|---|---|
| tamoxifen | L | 210 | 77 | 2,727272727 |
| thioguanine | L | 441,533 | 26400 | 0,016724735 |
| baricitinib | L | 2 | 2535 | 0,000788955 |
| diclofenac | M | 19796 | 0,05 | 395920 |
| ibuprofen | M | 104227 | 5 | 20845,4 |
| paracetamol | M | 1260000 | 500 | 2520 |
| naproxen | M | 287130 | 1700 | 168,9 |
| ketoprofen | M | 29000 | 1240 | 23,38709677 |
| flufenamic acid | M | 2509 | 400 | 6,2725 |
| mefenamic acid | M | 5416 | 1000 | 5,416 |
| meclofenamic acid | M | 285 | 97,26 | 2,930289944 |
| celecoxib | M | 186,3375 | 90 | 2,070416667 |
| niflumic acid | M | 587,73 | 791 | 0,743021492 |
| etodolac | M | 156,521 | 793 | 0,19737831 |
| indometacin | M | 82,707 | 3220 | 0,025685404 |
| piroxicam | M | 22,963 | 2209 | 0,010395201 |
| ketorolac | M | 14,829 | 2087 | 0,007105414 |
| methocarbamol | M | 142 | 24882,28 | 0,005706873 |
| alminoprofen | M | 5 | 1057 | 0,004730369 |
| 4-acetamidoantipyrine | metabolite | 27079 | 100 | 270,79 |
| anhydro erythromycin | metabolite | 7996 | 37 | 216,1081081 |
| o-desmethylvenlafaxine | metabolite | 5028 | 880 | 5,713636364 |
| venlafaxine-o-desmethyl | metabolite | 976 | 880 | 1,109090909 |
| lidocaine n-oxide | metabolite | 1432 | 1587 | 0,902331443 |
| 2-ethylidene-1-5-dimethyl-3,3-di | metabolite | 68 | 127,42 | 0,533668184 |
| venlafaxine-n-desmethyl | metabolite | 2278 | 6613,84 | 0,344429257 |
| metoprolol acid | metabolite | 630 | 1940 | 0,324742268 |
| methyl 3,4-dihydroxybenzoate | metabolite | 1379,259 | 6145 | 0,224452238 |
| cbz 10,11-epoxide | metabolite | 365,967 | 1879 | 0,194766897 |
| hydroxymetronidazole | metabolite | 523,53 | 3693 | 0,141762794 |
| venlafaxine-n,n-didesmethyl | metabolite | 1185 | 13980,36 | 0,084761766 |
| o-desmethyltramadol | metabolite | 670 | 9122,843026 | 0,073442018 |
| norlidocaine | metabolite | 89,9735 | 1514 | 0,059427675 |
| n-desmethyltramadol | metabolite | 495 | 10956,7 | 0,045177836 |
| 2-acetamidophenol | metabolite | 173,831 | 4483,13734 | 0,038774409 |
| 4-formylaminoantipyrine | metabolite | 38,388 | 1262 | 0,030418384 |
| albendazole-2-aminosulfone | metabolite | 25,996 | 877 | 0,029641961 |
| venlafaxine-n,o-didesmethyl | metabolite | 324 | 11393,95 | 0,028436144 |
| amisulpride n-oxide | metabolite | 42,797 | 2298 | 0,018623586 |
| n4-acetyl-sulfamethazine | metabolite | 0,9 | 236,21 | 0,003810169 |
| 8-methoxypsoralen | metabolite | 5 | 1843,01 | 0,002712953 |
| n4-acetyl-sulfamerazine | metabolite | 1,6 | 592,46 | 0,002700604 |
| venlafaxine n-oxide | metabolite | 2 | 2981 | 0,000670916 |
| n-desmethylvenlafaxine | metabolite | 2 | 3023 | 0,000661594 |
| n4-acetyl-sulfapyridine | metabolite | 1310 | | |
| cbz 10,11-trans-dixydroxy 10,11- | metabolite | 650 | | |
| 4'-hydroxydiclofenac | metabolite | 555 | | |
| paraxanthine | metabolite | 380,874 | | |
| clindamycin sulfoxide | metabolite | 220 | | |
| n4-acetylsulfapyridine | metabolite | 61 | | |

| | | | | |
|---|---|---|---|---|
| noroxicodone | metabolite | 54 | | |
| n4-acetyl-sulfamethoxazole | metabolite | 43 | | |
| n desmethyl citalopram | metabolite | 42 | | |
| cbz 10,11-dihydro | metabolite | 40 | | |
| hydroxybupropione | metabolite | 37 | | |
| n-desmethylclarithromycin | metabolite | 26 | | |
| n-bisdesmethyltramadol | metabolite | 11 | | |
| n4-acetyl-sulfadinazine | metabolite | 1,5 | | |
| carbamazepine | N | 10300 | 10 | 1030 |
| temazepam | N | 4504 | 7,08 | 636,1581921 |
| codeine | N | 20030 | 60 | 333,8333333 |
| citalopram | N | 2364,35 | 10 | 236,435 |
| tramadol | N | 6444 | 32 | 201,375 |
| venlafaxine | N | 8273 | 70 | 118,1857143 |
| lidocaine | N | 133910 | 2610 | 51,30651341 |
| sertraline | N | 417,57 | 9,4 | 44,42234043 |
| sucralose | N | 1100000 | 29694,27 | 37,04418395 |
| amitriptyline | N | 2232 | 135,76 | 16,44077784 |
| antipyrine | N | 3242 | 280 | 11,57857143 |
| lorazepam | N | 978 | 95,68 | 10,22157191 |
| clozapine | N | 356 | 37 | 9,621621622 |
| nicotine | N | 11700 | 1390 | 8,417266187 |
| morphine | N | 8200 | 1330 | 6,165413534 |
| fluoxetine | N | 370 | 100 | 3,7 |
| entacapone | N | 2900 | 854 | 3,395784543 |
| clonazepam | N | 53 | 19,39 | 2,733367715 |
| trazodone | N | 41 | 15,85 | 2,586750789 |
| diazepam | N | 490 | 291 | 1,683848797 |
| salicylic acid | N | 55086 | 37900 | 1,453456464 |
| hydrocodone | N | 1000 | 939 | 1,064962726 |
| methadone | N | 170 | 170 | 1 |
| lidocaine n-oxide | N | 1432 | 1587 | 0,902331443 |
| alprazolam | N | 56 | 76,84 | 0,72878709 |
| thioridazine | N | 485,31 | 734,818 | 0,660449254 |
| tiapride | N | 3313 | 5080 | 0,652165354 |
| nortriptylin | N | 101 | 190 | 0,531578947 |
| primidone | N | 3700 | 9113,83 | 0,405976412 |
| sulpiride | N | 1100 | 3595 | 0,305980529 |
| oxcarbazepine | N | 515,848 | 2117 | 0,243669343 |
| maprotiline | N | 37 | 158 | 0,234177215 |
| ketamine | N | 166 | 720 | 0,230555556 |
| bupropione | N | 194 | 950 | 0,204210526 |
| zaleplon | N | 8 | 46,04 | 0,173761946 |
| gabapentin | N | 16408 | 100000 | 0,16408 |
| propyphenazone | N | 115 | 800 | 0,14375 |
| amantadine | N | 3182 | 25000 | 0,12728 |
| lamotrigine | N | 991 | 8000 | 0,123875 |
| risperidone | N | 45 | 380 | 0,118421053 |
| memantine | N | 194 | 1840 | 0,105434783 |
| oxazepam | N | 36 | 370 | 0,097297297 |

| | | | | |
|---|---|---|---|---|
| meprobamate | N | 2500 | 26977,41 | 0,092670127 |
| benzatropine | N | 23 | 270 | 0,085185185 |
| pregabalin | N | 7680 | 100000 | 0,0768 |
| zolpidem | N | 12 | 177,52 | 0,067598017 |
| oxycodone | N | 460 | 8040 | 0,05721393 |
| orphenadrine | N | 79 | 1450 | 0,054482759 |
| rizatriptan | N | 20 | 400 | 0,05 |
| ropinirole | N | 29 | 623,97 | 0,046476593 |
| tacrine | N | 24 | 520 | 0,046153846 |
| tapentadol | N | 1923 | 53000 | 0,036283019 |
| mirtazapine | N | 36 | 1000 | 0,036 |
| methylphenidate | N | 202 | 11600 | 0,017413793 |
| topiramate | N | 1025 | 93000 | 0,011021505 |
| norfentanyl | N | 320 | 73000 | 0,004383562 |
| ethenzamide | N | 45,381 | 10400 | 0,004363558 |
| amisulpride | N | 600 | 140000 | 0,004285714 |
| phenytoin | N | 620 | 163000 | 0,003803681 |
| salicylamide | N | 50 | 16000 | 0,003125 |
| quetiapine | N | 19 | 10000 | 0,0019 |
| fentanyl | N | 0,307 | 171,35 | 0,001791655 |
| phenacetin | N | 56,286 | 41000 | 0,001372829 |
| phenobarbital | N | 4,95 | 4590,98 | 0,001078201 |
| haloperidol | N | 1,2 | 1400 | 0,000857143 |
| prilocaine | N | 92 | 150000 | 0,000613333 |
| diatrizoic acid | N | 292 | 10000000 | 0,0000292 |
| nordiazepam | N | 80 | | |
| albendazole | P | 10880 | 6,023156828 | 1806,361732 |
| permethrin | P | 100 | 0,811 | 123,3045623 |
| ivermectin | P | 1590 | 141,63 | 11,22643508 |
| hydroxychloroquine | P | 372 | 70,51 | 5,275847398 |
| fenbendazole | P | 340 | 234 | 1,452991453 |
| mebendazole | P | 128 | 159,38 | 0,803112059 |
| pyrantel | P | 166 | 2445,92 | 0,067868123 |
| levamisole | P | 81 | 1810 | 0,044751381 |
| praziquantel | P | 27,4545 | 1340 | 0,020488433 |
| thiabendazole | P | 107 | | |
| cetirizine | R | 9296,714 | 410,83 | 22,62910206 |
| diphenhydramine | R | 573,918 | 80 | 7,173975 |
| salbutamol | R | 2800 | 1158 | 2,417962003 |
| epinastine | R | 623,241 | 627 | 0,994004785 |
| levocabastine | R | 47 | 170 | 0,276470588 |
| clenbuterol | R | 9,9 | 166 | 0,059638554 |
| ketotifen | R | 10 | 254,84 | 0,039240308 |
| dextrorphan | R | 43 | 1360,65 | 0,031602543 |
| pheniramine | R | 25 | 1140,03 | 0,021929247 |
| promethazine | R | 5 | 355 | 0,014084507 |
| chlorpheniramine | R | 67 | 17800 | 0,003764045 |
| fexofenadine | R | 580 | 200000 | 0,0029 |
| doxylamine | R | 46 | | |
| benproperine | R | 31 | | |

| | | | | |
|---|---|---|---|---|
| loratadine | R | | | |
| cinchocaine | S | 14,503 | 2505,533945 | 0,005788387 |
| iopamidol | V | 570000 | 30,11 | 18930,58784 |
| iohexol | V | 12900 | 138,84 | 92,91270527 |
| iopromide | V | 783 | 10000000 | 0,0000783 |
| iomeprol | V | 801 | | |
| tilmycosin | VET | 420000 | 500 | 840 |
| tiamulin | VET | 21000 | 1000 | 21 |
| morantel | vet | 27000 | 1357,94 | 19,88305816 |
| marbofloxacin | VET | 6300 | 7780 | 0,809768638 |
| florfenicol | VET | 119,9495 | 1421,728708 | 0,084368768 |
| sulfanitran | VET | 31 | 885,97 | 0,034989898 |
| cyromazine | VET | 4,4045 | 2500 | 0,0017618 |
| ipronidazole | VET | 2 | 12300 | 0,000162602 |
| ronidazole | Vet | 3664 | | |

| risk_level |
|---|
| High |
| Moderate |
| Low |
| Low |
| Low |
| Low |
| Low |
| Low |
| Low |
| Insignificant |
| Insignificant |
| Insignificant |
| Insignificant |
| High |
| Low |
| Low |
| High |
| High |
| High |
| High |
| High |
| High |
| High |
| High |
| High |
| Moderate |
| Moderate |
| Moderate |
| Moderate |
| Moderate |
| Low |
| Low |
| Low |
| Low |
| Low |
| Low |
| Low |
| Low |
| Low |
| Low |
| Low |
| Low |
| Low |
| Low |
| Low |
| Low |
| Insignificant |
| Insignificant |
| Insignificant |

Insignificant
Insignificant
High
High
Moderate
Insignificant

High
High
High
High
Moderate
Moderate
Low
Low
Low
Low
Low
Insignificant

High
Low
Low
Insignificant
High
High
High
High
High
High
High
High
High
High
High
High
High
High
High
High
High
High
High
High
High
High
High
High
High
High

High
High
High
High
Moderate
Moderate
Moderate
Moderate
Moderate
Moderate
Moderate
Moderate
Moderate
Low
Low
Low
Low
Low
Low
Low
Low
Low
Low
Low
Low
Low
Low
Insignificant
Insignificant
Insignificant
Insignificant
Insignificant
Insignificant
Insignificant
Insignificant
Insignificant
Insignificant
Insignificant
Insignificant
Insignificant

High
High
High
High
High
Moderate

Moderate
Low
Insignificant
High
High
High
High
High
Moderate
Moderate
Moderate
Moderate
Low
Low
Low
Low
Insignificant
Insignificant
Insignificant
High
High
Moderate
Moderate
Low
Low
Low
Low
Low
Low
Low
Low
Low
Low
Low
Low
Low
Low
Low
Low
Insignificant
Insignificant
Insignificant
Insignificant
Insignificant

High
High
High
High
High
High
High
High
High
High
High
High
Moderate
Moderate
Moderate
Moderate
Moderate
Moderate
Moderate
Moderate
Moderate
Moderate
Low
Low
Low
Low
Low
Low
Low
Low
Low
Low
Low
Low
Low
Low
Low
Low
Low
Low
Low
Low

Low
Low
Low
Low
Low
Low
Low
Low
Low
Low
Low
Low
Low
Insignificant
Insignificant
Insignificant
Insignificant
Insignificant
Insignificant
Insignificant
Insignificant
Insignificant
Insignificant
Insignificant
Insignificant

High
High
High
Moderate
Moderate
Low
Low
Low
Low

High
Moderate
Moderate
Low
Low
Low
Low
Low
Low
Low
Insignificant
Insignificant

Insignificant
High
High
Insignificant

High
High
High
Low
Low
Low
Insignificant
Insignificant
