## Supplementary material for "The environmental impact of pharmaceuticals: an evidence-mapping review of recent data on aquatic concentrations and predictable effects": Table S1 Table S2 Table S3: S2 Supplementary Material - MEC.pdf

| API | MEC (ng/L | Type of val | Compartm | Location | Article |
|---|---|---|---|---|---|
| alminoprof | 5 | max | surface wa | South Kore | Kim et al. (2023) |
| celecoxib | 17 | max | Surface wa | UK, River T | Egli et al. (2023) |
| celecoxib | 186.3375 | max | Surface wa | South Kore | Kim et al. (2023) |
| celecoxib | 20 | max | Influent W | Spain, Bas | Lopez-Herguedas et al. (2023) |
| diclofenac | 1300 | max | Sea water | Turkey, Ma | Korkmaz et al. (2021) |
| diclofenac | 1000 | max | Sanitary O | Australia, S | Besley et al. (2023) |
| diclofenac | 10000 | max | Household | Finland | Äystö et al. (2023) |
| diclofenac | 1100 | max | wastewate | Czech Rep | Kodesova et al. (2024) |
| diclofenac | 1290 | max | Effluent W | Northern E | Spilsbury et al. (2024) |
| diclofenac | 1600 | avg | Surface wa | India, Beng | Nozaki et al. (2023) |
| diclofenac | 176.4725 | max | Surface wa | South Kore | Kim et al. (2023) |
| diclofenac | 18 | max | Surface wa | India, Gang | Richards et al. (2023) |
| diclofenac | 19796 | max | Influent W | North Port | Montes et al. (2023) |
| diclofenac | 2201.7 | max | Surface wa | Hungary, B | Staszny et al. (2021) |
| diclofenac | 515.3 | max | Surface wa | Ecuador, E | Cipriani-Avila et al. (2023) |
| diclofenac | 518 | max | Surface wa | Cameroon | Branchet et al. (2019) |
| diclofenac | 527 | max | Surface wa | UK, River T | Egli et al. (2023) |
| diclofenac | 7.1 | max | Surface wa | Antarctica | Postigo et al. (2023) |
| diclofenac | 746.9 | max | Wastewate | Antarctica, | Szopińska et al. (2021) |
| diclofenac | 922 | max | Influent W | Ireland, We | Rapp-Wright et al. (2023) |
| diclofenac | 970 | max | Influent W | Spain, Cad | Santana-Viera et al. (2023) |
| etodolac | 156.521 | max | Surface wa | South Kore | Kim et al. (2023) |
| etodolac | 33 | max | Influent W | Spain, Cad | Santana-Viera et al. (2023) |
| flufenamic | 2509 | max | Effluent W | North Port | Montes et al. (2023) |
| flufenamic | 737 | max | Influent W | Spain, Cad | Santana-Viera et al. (2023) |
| ibuprofen | 2130 | max | Sea water | Turkey, Ma | Korkmaz et al. (2021) |
| ibuprofen | 104227 | max | Influent W | Spain, Cad | Santana-Viera et al. (2023) |
| ibuprofen | 13000 | max | Sanitary O | Australia, S | Besley et al. (2023) |
| ibuprofen | 1800 | max | Surface wa | Spain, Tag | Royano et al. (2023) |
| ibuprofen | 2332 | max | Influent W | Indonesia | Astuti et al. (2022) |
| ibuprofen | 2400 | avg | Surface wa | India, Beng | Nozaki et al. (2023) |
| ibuprofen | 276 | max | Surface wa | Cameroon | Branchet et al. (2019) |
| ibuprofen | 476.61 | max | Wastewater | | Szopińska et al. (2021) |
| ibuprofen | 502 | max | Effluent W | Northern E | Spilsbury et al. (2024) |
| ibuprofen | 81 | max | Hospital w | Finland | Äystö et al. (2023) |
| ibuprofen | 89000 | max | Influent W | USA, South | Gerrity et al. (2023) |
| indometac | 34 | avg | Surface wa | India, Beng | Nozaki et al. (2023) |
| indometac | 61 | max | Effluent W | Spain, Bas | Lopez-Herguedas et al. (2023) |
| indometac | 82.707 | max | Surface wa | South Kore | Kim et al. (2023) |
| ketoprofen | 370 | max | Sea water | Turkey, Ma | Korkmaz et al. (2021) |
| ketoprofen | 14.61 | max | Wastewate | Antarctica, | Szopińska et al. (2021) |
| ketoprofen | 1479 | max | Influent W | Spain, Cad | Santana-Viera et al. (2023) |
| ketoprofen | 29000 | max | Hospital w | Finland | Äystö et al. (2023) |
| ketoprofen | 59.5 | max | Groundwa | Poland | Nikolenko et al. (2023) |
| ketoprofen | 607 | max | Surface wa | Spain, Tag | Royano et al. (2023) |
| ketoprofen | 780 | max | Effluent W | Spain, Bas | Lopez-Herguedas et al. (2023) |
| ketorolac | 14.829 | max | Surface wa | South Kore | Kim et al. (2023) |

| | | | | | |
|---|---|---|---|---|---|
| meclofena | 285 | max | Influent W\ | Spain, Cad | Santana-Viera et al. (2023) |
| mefenami | 1463 | max | Influent W\ | Ireland, W | Rapp-Wright et al. (2023) |
| mefenami | 166 | max | Surface wa | UK, River T | Egli et al. (2023) |
| mefenami | 2600 | avg | Surface wa | Indonesia, | Nozaki et al. (2023) |
| mefenami | 261.672 | max | surface wa | South Kore | Kim et al. (2023) |
| mefenami | 5416 | max | Hospital w | Tunisia, Tu | Nasri et al. (2024) |
| methocarb | 142 | max | Surface wa | USA, Poke | Pronschinske et al. (2023) |
| naproxen | 3960 | max | Surface wa | Pakistan, L | Wilkinson et al. (2022) |
| naproxen | 287130 | max | Surface wa | Hungary, B | Staszny et al. (2021) |
| naproxen | 1360 | max | hospital wa | Tunisia, So | Nasri et al. (2024) |
| naproxen | 25000 | max | Hospital w | Finland | Äystö et al. (2023) |
| naproxen | 2653.1 | max | Wastewate | Antarctica, | Szopińska et al. (2021) |
| naproxen | 31340 | max | influent W\ | Spain, Cad | Santana-Viera et al. (2023) |
| naproxen | 49000 | max | Influent W\ | USA, South | Gerrity et al. (2023) |
| naproxen | 554 | max | Surface wa | Spain, Tag | Royano et al. (2023) |
| naproxen | 906.768 | max | Surface wa | South Kore | Kim et al. (2023) |
| naproxen | 9640 | max | Effluent W\ | Spain, Bas | Lopez-Herguedas et al. (2023) |
| naproxen | 3000 | max | Sanitary O\ | Australia, S | Besley et al. (2023) |
| niflumic ac | 14 | max | Influent W\ | Spain, Cad | Santana-Viera et al. (2023) |
| niflumic ac | 587.73 | max | Surface wa | South Kore | Kim et al. (2023) |
| paracetam | 227000 | max | Surface wa | Bolivia, La | wilkinson et al. (2022) |
| paracetam | 115493 | max | Influent W\ | Spain, Cad | Santana-Viera et al. (2023) |
| paracetam | 1168 | max | Surface wa | Spain, Tag | Royano et al. (2023) |
| paracetam | 1260000 | max | hospital wa | Tunisia, Tu | Nasri et al. (2024) |
| paracetam | 169.8 | max | Surface wa | Ecuador, E | Cipriani-Avila et al. (2023) |
| paracetam | 181000 | max | Sanitary O\ | Australia, S | Besley et al. (2023) |
| paracetam | 1921.616 | max | Surface wa | South Kore | Kim et al. (2023) |
| paracetam | 227000 | max | Surface wa | Bolivia, La | wilkinson et al. (2022) |
| paracetam | 332.42 | max | Wastewate | Antarctica, | Szopińska et al. (2021) |
| paracetam | 45.8 | max | wastewate | Kenya, Nai | Bagnis et al. (2019) |
| paracetam | 450000 | max | hospital wa | Finland | Äystö et al. (2023) |
| paracetam | 550000 | max | Influent W\ | USA, South | Gerrity et al. (2023) |
| paracetam | 550820 | max | Surface wa | Hungary, B | Staszny et al. (2021) |
| paracetam | 5660 | max | Surface wa | Cameroon | Branchet et al. (2019) |
| paracetam | 67564.102 | max | Influent W\ | North Port | Montes et al. (2023) |
| paracetam | 80.5 | max | Surface wa | USA, Poke | Pronschinske et al. (2023) |
| paracetam | 85774 | max | Effluent W\ | Spain, Bas | Lopez-Herguedas et al. (2023) |
| paracetam | 9248 | max | Influent W\ | Indonesia | Astuti et al. (2022) |
| piroxicam | 22.963 | max | Surface wa | South Kore | Kim et al. (2023) |
