## Supplementary material for "The environmental impact of pharmaceuticals: an evidence-mapping review of recent data on aquatic concentrations and predictable effects": Table S1 Table S2 Table S3: S3 Supplementary Material - PNEC.pdf

| API | PNEC (ng/L) | Species | AF | Article |
|---|---|---|---|---|
| 5-fluorocytosine | 0,22168 | Algae | 1000 | Santana-Viera et al. (2023) |
| amoxicillin | 78 | Algae | 10 | Dulio et al. (2020) |
| ampicillin | 12 | | | Kim et al. (2023) |
| atazanavir | 1771 | | | Kim et al. (2023) |
| azithromycin | 19 | Fish | 1000 | Dulio et al. (2020) |
| azithromycin | 19 | | | Montes et al. (2023) |
| azithromycin | 19 | | | Kim et al. (2023) |
| azithromycin | 19 | | | Egli et al. (2023) |
| azithromycin | 1874 | Algae | | Li et al. (2023) |
| cefadroxil | 2670 | | | Kim et al. (2023) |
| cefradine | 5893,697436 | Algae | 1000 | Dulio et al. (2020) |
| cephalexin | 4026,24 | Algae | 1000 | Dulio et al. (2020) |
| chloramphenicol | 130 | Algae | 1000 | Lei K. et al. (2019) |
| chloramphenicol | 198,7690299 | | | Kim et al. (2023) |
| chloramphenicol | 2728,71 | Algae | | Dulio et al. (2020) |
| chloramphenicol | 2730 | | | Egli et al. (2023) |
| chlortetracycline | 500 | | | Watanabe et al. (2023) |
| ciprofloxacin | 5 | Algae | | Li et al. (2023) |
| ciprofloxacin | 5 | Algae | 1000 | Äystö et al. (2023) |
| ciprofloxacin | 60 | | | Watanabe et al. (2023) |
| ciprofloxacin | 89 | Algae | 50 | Dulio et al. (2020) |
| ciprofloxacin | 110 | Daphnia | | Liu et al. (2023) |
| ciprofloxacin | 470 | Algae | 10 | Bouzas-Monroy et al. (2022) |
| ciprofloxacin | 1100 | Crustaceans | | Lei K. et al. (2019) |
| clarithromycin | 120 | Algae | 10 | Dulio et al. (2020) |
| clarithromycin | 120 | | | Montes et al. (2023) |
| clarithromycin | 120 | | | Kim et al. (2023) |
| clarithromycin | 120 | | | Egli et al. (2023) |
| clarithromycin | 245 | algae | 10 | Nozaki et al. (2023) |
| clarithromycin | 250 | Algae | 10 | Bouzas-Monroy et al. (2022) |
| clarithromycin | 1400 | Algae | | Liu et al. (2023) |
| clevudine | 1592 | | | Kim et al. (2023) |
| clindamycin | 1900 | Fish | | Spilsbury et al. (2024) |
| danofloxacin | 93,08 | Fish | 1000 | Dulio et al. (2020) |
| darunavir | 1815 | | | Kim et al. (2023) |
| difloxacin | 240000 | | 1000 | Zhang F et al. (2023) |
| doxycycline | 2000 | | | Watanabe et al. (2023) |
| efavirenz | 200,79 | Fish | 1000 | Dulio et al. (2020) |
| enoxacin | 2514,2 | Algae | 1000 | Dulio et al. (2020) |
| enrofloxacin | 60 | | | Watanabe et al. (2023) |
| erythromycin | 20 | Algae | | Lei K. et al. (2019) |
| erythromycin | 200 | Algae | 10 | Bouzas-Monroy et al. (2022) |
| erythromycin | 200 | algae | 10 | Nozaki et al. (2023) |
| erythromycin | 300 | | | Dulio et al. (2020) |
| erythromycin | 500 | | | Watanabe et al. (2023) |
| erythromycin | 940 | Daphnia | | Liu et al. (2023) |

| | | | | |
|---|---|---|---|---|
| fleroxacin | 1112,28 | Fish | 1000 | Dulio et al. (2020) |
| flubendazole | 7,90569415 | | | Kim et al. (2023) |
| flubendazole | 235,94 | Algae | 1000 | Dulio et al. (2020) |
| fluconazole | 250 | | | Bagnis et al. (2019) |
| fluconazole | 1043,31 | Algae | 1000 | Dulio et al. (2020) |
| fluconazole | 3060 | | | Kim et al. (2023) |
| fluconazole | 50000 | Fish | 1000 | Bouzas-Monroy et al. (2022) |
| flumequine | 1500 | | | Kim et al. (2023) |
| itraconazole | 12,88 | Algae | 1000 | Dulio et al. (2020) |
| lincomycin | 14 | | | Kim et al. (2023) |
| lincomycin | 70 | Algae | | Liu et al. (2023) |
| lincomycin | 780 | algae | 10 | Nozaki et al. (2023) |
| lincomycin | 810 | | | Watanabe et al. (2023) |
| lincomycin | 3948,62 | Algae | 1000 | Dulio et al. (2020) |
| lincomycin | 3950 | | | Egli et al. (2023) |
| lincomycin | 7800 | Algae | 10 | Bouzas-Monroy et al. (2022) |
| lomefloxacin | 826,18 | Fish | 1000 | Dulio et al. (2020) |
| lopinavir | 1575 | | | Kim et al. (2023) |
| marbofloxacin | 7780 | Algae | 1000 | Dulio et al. (2020) |
| metacycline | 6270 | Algae | | Liu et al. (2023) |
| metronidazole | 130 | | | Bagnis et al. (2019) |
| metronidazole | 33081,17 | Algae | 1000 | Dulio et al. (2020) |
| metronidazole | 86053,64534 | | | Kim et al. (2023) |
| metronidazole | 100000 | Fish | 1000 | Daniele et al. (2023) |
| nevirapine | 481,82 | Algae | 1000 | Dulio et al. (2020) |
| nevirapine | 43000 | Algae | 1000 | Bouzas-Monroy et al. (2022) |
| norfloxacin | 500 | | | Watanabe et al. (2023) |
| ofloxacin | 21 | Algae | | Liu et al. (2023) |
| ofloxacin | 21 | Algae | | Lei K. et al. (2019) |
| ofloxacin | 22 | Bacteria | 1000 | Äystö et al. (2023) |
| ofloxacin | 1387,89 | Fish | 1000 | Dulio et al. (2020) |
| ofloxacin | 1440 | Algae | | Li et al. (2023) |
| orbifloxacin | 500 | | | Watanabe et al. (2023) |
| oseltamivir | 3352 | | | Kim et al. (2023) |
| oseltamivir | 13022,98 | Algae | 1000 | Dulio et al. (2020) |
| oseltamivir | 1000000 | Algae | 10 | Bouzas-Monroy et al. (2022) |
| oxolinic acid | 380 | | | Kim et al. (2023) |
| oxytetracycline | 500 | | | Watanabe et al. (2023) |
| pefloxacin | 2575,27 | Algae | 1000 | Dulio et al. (2020) |
| penicillin | 2135,24 | Algae | 1000 | Dulio et al. (2020) |
| penicillin v | 180,681 | Algae | | Dulio et al. (2020) |
| ritonavir | 4,61 | Algae | | Dulio et al. (2020) |
| roxithromycin | 100 | Algae | | Lei K. et al. (2019) |
| roxithromycin | 100 | algae | 10 | Nozaki et al. (2023) |
| roxithromycin | 4660 | Algae | | Liu et al. (2023) |
| sofosbuvir | 3117 | | | Kim et al. (2023) |
| streptomycin | 482,8664412 | | | Kim et al. (2023) |
| sulfabenzamide | 3052,62 | Algae | | Dulio et al. (2020) |

| | | | | |
|---|---|---|---|---|
| sulfacetamide | 14307,8 | Algae | | Dulio et al. (2020) |
| sulfachlopyridazine | 800 | Daphnia | | Liu et al. (2023) |
| sulfadiazine | 100 | Algae | | Lei K. et al. (2019) |
| sulfadiazine | 1000 | Daphnia | | Liu et al. (2023) |
| sulfadiazine | 2200 | algae | | Spilsbury et al. (2024) |
| sulfadiazine | 13000 | Algae | 10 | Bouzas-Monroy et al. (2022) |
| sulfadimethoxine | 600 | | | Watanabe et al. (2023) |
| sulfadimethoxine | 2259 | Daphnia | | Liu et al. (2023) |
| sulfamerazine | 780 | algae | 100 | Nozaki et al. (2023) |
| sulfamethazine | 700 | Daphnia | | Liu et al. (2023) |
| sulfamethazine | 63000 | algae | 10 | Nozaki et al. (2023) |
| sulfamethizole | 1495,22 | Algae | | Dulio et al. (2020) |
| sulfamethoxazole | 150 | Algae | | Liu et al. (2023) |
| sulfamethoxazole | 210 | Daphnia | | Li et al. (2023) |
| sulfamethoxazole | 560 | | | Bagnis et al. (2019) |
| sulfamethoxazole | 590 | Algae | 10 | Bouzas-Monroy et al. (2022) |
| sulfamethoxazole | 590 | algae | 10 | Nozaki et al. (2023) |
| sulfamethoxazole | 600 | | | Kim et al. (2023) |
| sulfamethoxazole | 600 | | | Watanabe et al. (2023) |
| sulfamethoxazole | 600 | | | Egli et al. (2023) |
| sulfamethoxypiridazine | 3820 | Algae | | Li et al. (2023) |
| sulfametoxydiazine | 800 | Daphnia | | Liu et al. (2023) |
| sulfamonomethoxine | 600 | | | Watanabe et al. (2023) |
| sulfamonomethoxine | 2085 | Daphnia | | Liu et al. (2023) |
| sulfanoxaline | 19700 | Algae | | Li et al. (2023) |
| sulfapyridine | 460 | | | Kim et al. (2023) |
| sulfapyridine | 460 | | | Egli et al. (2023) |
| sulfapyridine | 700 | Daphnia | | Liu et al. (2023) |
| sulfaquinoxaline | 143,05 | Algae | | Dulio et al. (2020) |
| sulfathiazole | 11000 | | | Kim et al. (2023) |
| tetracycline | 1000 | | | Watanabe et al. (2023) |
| tetracycline | 3200 | Algae | 10 | Bouzas-Monroy et al. (2022) |
| trimethoprim | 5,8 | Mollusks | | Lei K. et al. (2019) |
| trimethoprim | 290 | Bacteria | 10 | Nozaki et al. (2023) |
| trimethoprim | 500 | | | Bagnis et al. (2019) |
| trimethoprim | 500 | | | Watanabe et al. (2023) |
| trimethoprim | 16000 | Algae | | Liu et al. (2023) |
| trimethoprim | 120000 | Algae | 10 | Dulio et al. (2020) |
| trimethoprim | 120000 | | | Kim et al. (2023) |
| trimethoprim | 120000 | | | Egli et al. (2023) |
| trimethoprim | 310000 | Algae | 10 | Bouzas-Monroy et al. (2022) |
| tylosin | 1000 | | | Watanabe et al. (2023) |
